# G-domain prediction across the diversity of G protein families

**DOI:** 10.1101/2019.12.24.888222

**Authors:** Hiral M. Sanghavi, Richa Rashmi, Anirban Dasgupta, Sharmistha Majumdar

## Abstract

Guanine nucleotide binding proteins are characterized by a structurally and mechanistically conserved GTP-binding domain, indispensable for binding GTP. The G domain comprises of five adjacent consensus motifs called G boxes, which are separated by amino acid spacers of different lengths. Several G proteins, discovered over time, are characterized by diverse function and sequence. This sequence diversity is also observed in the G box motifs (specifically the G5 box) as well as the inter-G box spacer length. The Spacers and Mismatch Algorithm (SMA) introduced in this study, can predict G-domains in a given G protein sequence, based on user-specified constraints for approximate G-box patterns and inter-box gaps in each G protein family. The SMA parameters can be customized as more G proteins are discovered and characterized structurally. Family-specific G box motifs including the less characterized G5 motif as well as G domain boundaries were predicted with higher precision. Overall, our analysis suggests the possible classification of G protein families based on family-specific G box sequences and lengths of inter-G box spacers.

**Significance Statement:** It is difficult to define the boundaries of a G domain as well as predict G boxes and important GTP-binding residues of a G protein, if structural information is not available. Sequence alignment and phylogenetic methods are often unsuccessful, given the sequence diversity across G protein families. SMA is a unique method which uses approximate pattern matching as well as inter-motif separation constraints to predict the locations of G-boxes. It is able to predict all G boxes including the less characterized G5 motif which marks the carboxy-terminal boundary of a G domain. Thus, SMA can be used to predict G domain boundaries within a large multi-domain protein as long as the user-specified constraints are satisfied.

**Classification:** Biological Sciences/Biophysics and Computational Biology

## Introduction

GTP binding proteins (G proteins) or GTPases, are known to play important roles in different cellular processes such as translation, cell signaling, nuclear and cytoplasmic protein transport, DNA repair and cytoskeleton stabilization (1, 2). GTPases are part of a larger family of NTPases (Nucleotide triphosphatase), which are proteins that bind and hydrolyze nucleotide triphosphates. NTPases are classified according to the presence of distinct structural folds; for example, the Rossman fold and the related FtsZ /tubulin fold, the P loop fold, the protein kinase fold, the HSP90/TopoII fold and the HSP70/RNaseH fold (3). Structurally the P loop is formed by an underlying consensus GXXXXGKS/T signature motif which folds into a pocket (3).

Most P loop GTPases, except for recently identified proteins like NACHT and McrB, are classified into two classes namely TRAFAC and SIMIBI, based on distinct sequence and structural signatures. The TRAFAC class includes translation factors, the Ras superfamily, Dynamin superfamily, heterotrimeric G proteins and Septins. The SIMIBI class includes Signal Recognition particle (SRP) GTPases, MinD and BioD related proteins. MinD and BioD related proteins lack specificity to bind GTP (3).

G proteins have a conserved “G domain” spanning 160-180 amino acid residues and with a molecular weight of 18-20 kDa. The G domain has the ability to bind both GTP and GDP; this allows the G protein to switch between a GTP-bound active state and a GDP-bound inactive state (when GTP is hydrolyzed to GDP). The G domain structurally folds into five α helices, six β strands and five loops interconnecting the helices and strands (1). Interestingly, the five loops within the G domain are involved in ligand (GTP) binding (1) (4) unlike other proteins where ligand binding sites are often alpha helical (5) or made of beta sheets (6). Each conserved loop, known as a G box, has a consensus amino acid sequence (7): (i) G1 box (P-loop/Walker A motif): “GXXXXGKS/T”. (ii) G2 box (switch 1/ effector domain) has a conserved Thr residue (iii) G3 box (Walker B motif/Switch II): “DXXG” (iv) G4 box: “NKXD” (v) G5 box: “EXSAX/ SAX”.

The crystal structures of G proteins with bound GTP/GDP/GTP analogues illustrate the importance of the most conserved amino acid residues in each G box of the G domain. Lys, a strongly conserved residue in the G1 box, the side chain of which directly interacts with the γ-phosphate oxygens of the bound nucleotide, is crucial for nucleotide binding. The conserved Ser of G1 box, Thr of the G2 box and Asp of the G3 box, do not directly contact the nucleotide but coordinate Mg2+, which bridges the β and γ phosphates of GTP. The Gly in the consensus G3 sequence DXXG is more variable than Asp. In the G4 box, the Asn contacts the C5 of purine base and the Asp forms a bifurcated hydrogen bond with the nitrogen atoms of the guanine base, thus conferring specificity to the guanidinium base (1). Although the G5 box is not well-characterized, structural evidence suggests that the Ala of the G5 box interacts with O6 of guanine base (1, 8) (Suppl. Figure 1).

The G boxes are separated from each other by amino acids (spacers) that are not involved in GTP binding. The spacer between G1 and G3 boxes is 40-80 residues long in most small G proteins (Ras superfamily) and 130-170 residues long in most large G proteins (Translation factor family), while the spacers between G3 and G4 boxes is between 40-80 residues long in most G proteins (7). It is difficult to predict a consensus spacer length between G1 and G2 since the G2 box consists of only one amino acid (Thr). The consensus spacer between G4 and G5 is unknown due to less available structural information about the G5 box.

Since the discovery of G proteins over fifty years ago, many new GTP binding proteins have been identified. These range from extensively studied proteins like Ras, Rab, Ran, Rac, Roc, Arf, translational factors and G***α*** subunit of heterotrimeric G proteins to more recently identified members like EngB, Septins, FeoB, AIG-1, OBG etc. These G proteins have very distinct cellular functions. For example, Ras is mainly involved in regulation of cell proliferation, differentiation, morphology and apoptosis. Ran plays an important role in nuclear-cytoplasmic transport, Rho/Rac is implicated in regulation of actin cytoskeletal organization and cell polarity while Rab proteins are involved in intracellular vesicular cargo transport. Arf proteins play an important role in vesicular protein trafficking via coat proteins and lipid modifying enzymes (9). Heterotrimeric G proteins are involved in signal transduction from cell surface G protein coupled receptors (GPCRs) to intracellular effectors (10). FeoB proteins, present only in archaeal and bacterial kingdoms, are found to transport ferrous ions in a potassium dependent manner and have an a GDI (GDP dissociation inhibitor) domain in addition to an amino-terminal G domain (11). Septins are of fundamental importance in cytokinesis, vesicular trafficking, phagocytosis and dendrite formation (12). AIG-1 is involved in T cell development and survival in animals and in response to biotic and abiotic stress in plants (13). Dynamins are implicated in budding of clathrin coated vesicles from the plasma membrane, mitochondrial division and interferon induced GTPase activity (14) while TrmE proteins are involved in tRNA wobble and uridine modification (15). OBG proteins are postulated to play important roles in many cellular processes like ribosome biogenesis and maturation, chromosome partitioning, DNA replication control (16). IRG (Immunity Related GTPases) are involved in immunity against pathogens in vertebrates (17). The signal recognition particle (SRP) along with its receptor constitutes essential cellular machinery that couples the synthesis of nascent proteins to their proper membrane localization (18). The Era family of proteins are speculated to be involved in translocation based on their association with 16s rRNA (19) and EngB proteins are involved in coordination of cell cycle events and assembly of 50S ribosome units in bacteria (20).The EngA/Der/YfgK/YphC class of proteins, which are yet to be structurally and functionally characterized, have the unique feature of two tandemly occurring GTPase domains separated by an acidic linker (20). Moreover, some G proteins like Septins, Dynamin, TrmE, IRG, AIG1/Toc are also known as GADs (GTPases activated by Dimerization) since their GTPase cycle is regulated by nucleotide binding and subsequent dimerization (21) while HflX class of proteins bind both ATP and GTP (22).

As more and more G proteins with diverse functions are identified and their corresponding structures are elucidated, it is important to investigate if all of these GTP-binding proteins, in fact, have a canonical G-domain with conserved G boxes and spacers, as described above. It is to be noted that there are many G proteins in which the conserved residues in each G box are substituted by other amino acids which do not abrogate nucleotide binding. For example, HypB (23) has Glu in place of Asp in the G3 box, YchF (24) has Glu in place of Asp in the G4 box, Septin family of proteins (25) have Ala or Gly in place of Asn in the G4 box and FeoB (26) and Ysxc, in which the conserved Ala in the G5 box, is substituted by Thr and Ser respectively (27, 28).

There are many freely available tools that can find motifs in sequences like ELM (Eukaryotic Linear Motif) (29), GTP binder (30), ScanProsite (31), Pfam (32) etc. ELM looks for short linear motifs (4-10 amino acids) in eukaryotic protein sequences; hence, given a G protein, ELM can possibly look for one G-Box at a time but not the entire G-domain. GTPbinder predicts GTP interacting dipeptides and tripeptides in a given protein sequence, using a machine learning based method; it uses PSSM which can consider possible variations in the sequence but cannot look for a G-Box (which is mostly longer) or the whole G-domain. Pfam identifies domains (in a submitted input protein sequence) that matches an already characterized protein domain and hence is not a good option for less studied proteins with very little domain information. ScanProsite is a more flexible method, which can look for motifs in a protein sequence as well as variations of the same; however, it is unable to look for different ranges of inter-motif gaps or spacers.

Due to the diversity in G box sequences and inter-G box spacer length, it is difficult to predict G boxes and/or important GTP-binding residues as well as define the boundaries of a G domain in a G protein, if structural information is not available. This task becomes more challenging in larger G proteins, like Translation factors, Septin, AIG1, HypB, which have multiple domains in addition to the G domain. In this study, we describe a method (“Spacers and Mismatch Algorithm” or SMA) that identifies all possible G domains in a given G protein sequence, after taking into account the G protein family-specific diversity. An exhaustive analysis of individual members of each of twenty different PROSITE G protein families was used to compile a set of approximate patterns for the G-boxes as well as a set of constraints for the inter-motif gaps. All of this information was then used by SMA to make its predictions. The novelty in our approach is that we use both approximate pattern matching as well as inter-motif separation constraints to predict the locations of the G-boxes. Our algorithm can be further customized to accommodate new information about mismatches (any amino acid) at any position of the G box as well as variations in the inter G box spacer length.

G box sequences predicted by SMA, were validated in corresponding G protein structures by observing if the predicted G boxes and G-domain to which they belonged, were indeed involved in guanine nucleotide binding. SMA was able to predict G protein family-specific G box consensus sequences for all G-boxes including the less characterized G5 Box. An accurate prediction of G5 box is specifically important since it is the last of five motifs in a G domain and hence marks the carboxy-terminal boundary of the domain. Thus, SMA analysis demonstrated that G protein families could possibly be classified based on family-specific G box sequences and inter-G box spacer lengths. Moreover, this method can be used to better predict the boundaries of a G domain, within a large multi-domain protein.

## Results

### Significant reduction in the size of protein families after removal of similarity bias

Twenty different G protein families (with a total of 8848 proteins across all families) were selected based on PROSITE documentation. Members of the same G protein family share very high sequence similarity amongst each other. In order to reduce redundancy in the results and over-representation of any sequence, the similarity bias from each G protein family was removed. As seen in Suppl Table 1, a significant reduction in the number of member proteins was observed in Arf, EngA, EngB, Era, OBG, Rac, Ran, Rab, Ras, SRP, translational and TrmE families. After removal of similarity bias, a total of 6487 proteins across all families were used for further analysis.

### Multiple Sequence alignment based phylogenetic relatedness of different G proteins

Proteins with high sequence identity often have similar functions and share evolutionary relationships amongst themselves. Thus, phylogenetic trees, constructed using Multiple Sequence Alignments (MSA) of protein sequences, group similar sequences together (33).

The Ras superfamily which includes Ras, Rac, Rab, Ran and Arf families, are small G proteins with a single G domain. However, the phylogenetic tree for the Ras superfamily demonstrates a distinct separation between individual families (9, 34). This indicates that there is a significant difference even amongst the members of the Ras superfamily. It was thus speculated that members of other G protein families will also be different from Ras superfamily members.

Phylogenetic trees, using Multiple Sequence Alignments (MSA) of full-length protein sequences, were constructed for each of the twenty G protein families. Some divergence was seen amongst the sequences of each G protein family. For example, in the Ras family, the human and mouse homologs of the protein AGAP3 group are close to each other but appear distant to the Ras proteins of different species like human, mouse, rat etc. (Suppl. File 2) However, in Era family, the human ERA protein clusters with the mouse, chicken and cow homologs but clusters away from the *Oryza sativa* ERA protein (Suppl. File 2)

In the translational family, all EF-Tu proteins cluster together, although within that cluster, human and mouse homologs of EF-Tu appear closer than that from *Thermotoga maritima* (Suppl. File 2). Different clustering patterns were thus seen in the phylogenetic trees generated for each G protein family.

Five representative proteins from each G protein family were arbitrarily chosen for phylogenetic analysis of G proteins across all twenty families (Figure 1). Only 5 members were chosen since it would be difficult to construct and visualise a phylogenetic tree with all the protein members of all twenty G protein families. Figure 1 illustrates that proteins that belong to the same family mostly cluster together for all the G protein families except for Roc, AIG1, translational, GB1/RHD3 and IRG G protein families.

**Figure 1.**
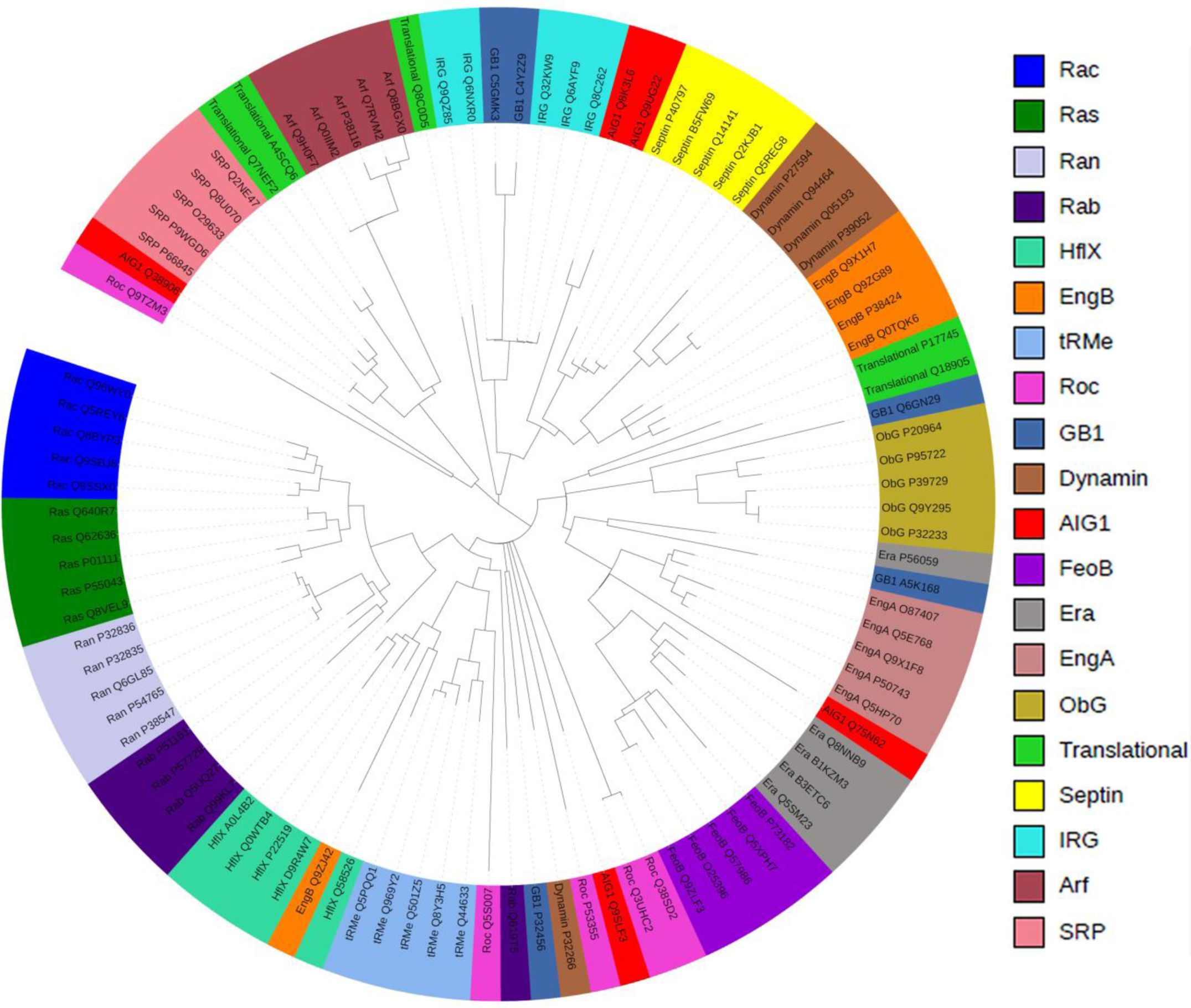
Phylogenetic trees of the twenty G protein families. Five proteins from each of the twenty G protein families were randomly selected to generate a phylogenetic tree. Tree scale 1.

It is to be noted, however, that the phylogenetic trees were constructed using full length protein sequences of individual G protein family members. Thus, the observed differences in Figure 1 may be explained by the fact that most of the G proteins, except the Ras superfamily members, are characterised by multi-domain proteins (one of these domains being the conserved G domain).

### Individual G boxes of the G domain cannot always be identified by Multiple Sequence Alignment

The GTP-binding domain (G domain) of a G protein can be defined by five G boxes separated by distinct spacings and with consensus sequences, namely G1-GXXXXGK, G3-DXXG, G4-NKXD, G5-SAX (where “X” =any amino acid). To investigate if the differences amongst members of different G protein families, observed in phylogenetic analysis (Figure 1) was due to differences in their individual G domains, we decided to look at their corresponding G domains more closely by performing multiple sequence alignment (MSA). However, MSA of all the proteins per G protein family (after removal of similarity bias) in each of the twenty G protein families did not align all the conserved residues of individual G boxes (for example, D and G in G3 box DXXG; Suppl. File 3) of a G domain.

G boxes could only be predicted upon careful visual examination of the alignment results i.e manually correlating the occurrence of conserved amino acids and spacers (For example, adjacent Gly and Lys residues occurring four amino acids downstream of a Gly at the amino-terminal end of Ras proteins can be predicted as the G1 box; a Gly residue occurring two residues downstream of an Asp residue, which again is separated from the predicted upstream G1 box by some spacer residues, is G3 box; and so on). The tedious manual method described above could predict G1, G3, G4 and G5 boxes only in Ras, Rab and Ran G protein families; G1, G3 and G4 boxes for Rho, Roc, Arf, Dynamin, EngA, EngB, FeoB, HflX, IRG, OBG, Septin, SRP and TrmE families; only G1 and G3 boxes in GB1/RHD3 and AIG1 families and no G box in the Translational family.

The difficulty in predicting consensus G boxes and their corresponding G domains in both the Ras family, with very similar protein sequences, to the Translational protein family, with very dissimilar protein sequences, maybe due to deletions, insertions or substitutions in otherwise similar protein sequences of the same G protein family. This could lead to a wide diversity of G box sequences as well as lengths of intervening spacers in the G domains of individual G protein families (35). Thus, although all G proteins are traditionally defined by a conserved G domain, there may be family-specific signatures for G box sequence and spacer length.

### Curating available information about the G protein families

To further probe our hypothesis that the diversity of G domains may have family-specificity, we embarked on an exhaustive analysis of all known G protein family members. After removal of similarity bias from the sequences of the twenty selected G-Protein families (Suppl. Table 1), G box consensus sequence and family specific spacers between consecutive G boxes were manually curated from the available literature and PDB database as described below:

#### Inter-G box Spacing

Structural information (obtained using X-ray crystallography or solution NMR) of G proteins bound to GTP or GTP analogues, can help directly determine their GTP binding sites and corresponding G domain. A few proteins from each G protein family, which have available structural information, were selected (Suppl. Table 2). The G domain of each of these proteins was mapped to the amino acid residues in the respective protein. Briefly, individual G box sequences were manually assigned to the amino acids that made contacts with either the phosphate (G1 and G3) or the guanine ring (G4 and G5) of GDP/GTP. The spacings between the consecutive G boxes was calculated by subtracting the amino acid position at the beginning of a specific G box from the amino acid position at the beginning of the succeeding G box (for example, if G1 box of a G protein starts at position 15 and the G3 box of the same protein starts at position 81, the spacing between them is calculated as 66). If there was structural information for more than one protein in a family, a putative range was decided based on the minimum and maximum values of calculated spacings for all such proteins within the same family (Suppl. Table 3). Some flexibility was allowed in these putative ranges. For instance, in well-studied families like Ras family which have shorter proteins, the putative spacer length was defined as +1 residues longer than the calculated spacer values, whereas for the translational or dynamin family which have longer proteins, the spacer length was defined as +2 residues longer than the calculated spacer boundaries. In less-studied protein families like AIG1, EngA which have short proteins, we have defined the spacer as +5 residues than the calculated spacer boundaries whereas for less studied protein families with long proteins like Septin and OBG, the spacer value was defined as +7 than the calculated spacer boundaries (Suppl. Table 3).

In some families like dynamin, EngB, HflX, OBG and translational, different proteins within the same family had significantly different inter G box spacers (Suppl. Table 4). Thus, for such families, more than one spacer constraint was included (Suppl. Table 3) to account for the observed intra-family variability.

In this way, putative spacings were defined for the approximate distances between G1-G3, G3-G4 and G4-G5 boxes (Suppl. Table 3) for each G protein family. Thus, we identified more precise spacings between consecutive G boxes in each G protein family as compared to the broad range of 40-80 amino acids between G1 box and G3 box and in between G3 box and G4 box suggested previously (7). Also, it was noted that the G4-G5 spacer length varied considerably and could be as short as 20 residues and as long as 120 residues.

#### Mismatch in G Box consensus sequence

Most G proteins (with structural information) have highly conserved G box sequences and do not differ from the consensus (i.e G1-GXXXXGK, G3-DXXG, G4-NKXD, G5-SAX; where “X” =any amino acid) by more than one mismatch.

For example, “GGAGIGF” (in Rbg1) and “AHIDAGK” (in S. *aureus* EF2) are G1 box motifs with one mismatch from the consensus motif, GXXXXGK. Also, “DIWE”, the G3 motif for human Rad and mouse REM2, has one mismatch from the consensus DXXG.

Most experimentally identified G4 boxes had 1 mismatch from the consensus sequence, NKXD [e.g., “TKLD” in human DNM1L (3W6N), “TQID” in human Cdc42 (1A4R), “TLRD” in human GBP1 (1DG3); “NVNE” in H. influenza YchF (1JAL);”G/AKXD” in human SEPT2 (2QAG), “T/CKXD” in Rac proteins, “TKXD” in EngB, Roc, SRP, Dynamin family], except the G4 boxes in the Roc, AIG-1, GB1/RHD3 and OBG families which usually had two mismatches [e.g., human LRRK2 “THLD” (2ZEJ), human GIMAP7 “RKEE” (3ZJC), human GBP1 “TLRD” (1DG3), human OLA1 “NLSE” (2OHF)].

It was observed that there were many variations to the G5 box consensus sequence”SAX”. This ranged from one mismatch in “SSE” in *B. subtilis* YsxC, “QAH”in *Legionella pneumophila* YsxC, “SSX” in EngB family, “SG/QX” in many translational and dynamin family proteins, “SCX/ CAX” in the Arf family, SGL” (4ZKD), “TAL” (2H5E), “SSK” (4U5X), “SCA” (1E0S), “SSA” (1MOZ), “NAT” (2ZEJ), “SQL” (3W6N), “STR” (3HYR) to “GVG” (2OG2), “GEG” (1OKK), “GVVRNSQ” in the dynamin family and”GXG/K” in SRP family. Thus, most identified G5 boxes, except GVG, GEG and GVVNRSQ, appeared to have one mismatch from the “SAX” consensus.

Therefore, based on our observed variations in G-box sequences, we defined that upto one mismatch was allowed in the consensus patterns for G1box (GXXXXGK), G3 box (DXXG), G4 (NKXD) and G5 box (SAX) for each of the G protein families. The only exception was the allowance of two mismatches in the G4 boxes of the Roc, AIG-1, GB1/RHD3 and OBG families.

A doughnut plot (Figure 2) and Suppl. Table 3 represent the manually curated information for all members across all G protein families. In Figure 2, each concentric circle represents consensus sequences of different G boxes, The longer the arc, higher is the occurrence of that consensus sequence across most G protein families; for example, NKXD is the most prevalent (occurs in 55% of curated G proteins) G4 box (green circle) while SAX is the most prevalent (occurs in 40% of curated G proteins) G5 box (blue circle). Figure 2 also shows that in up to 40% of curated G proteins, the G5 box sequence was unknown.

**Figure 2.**
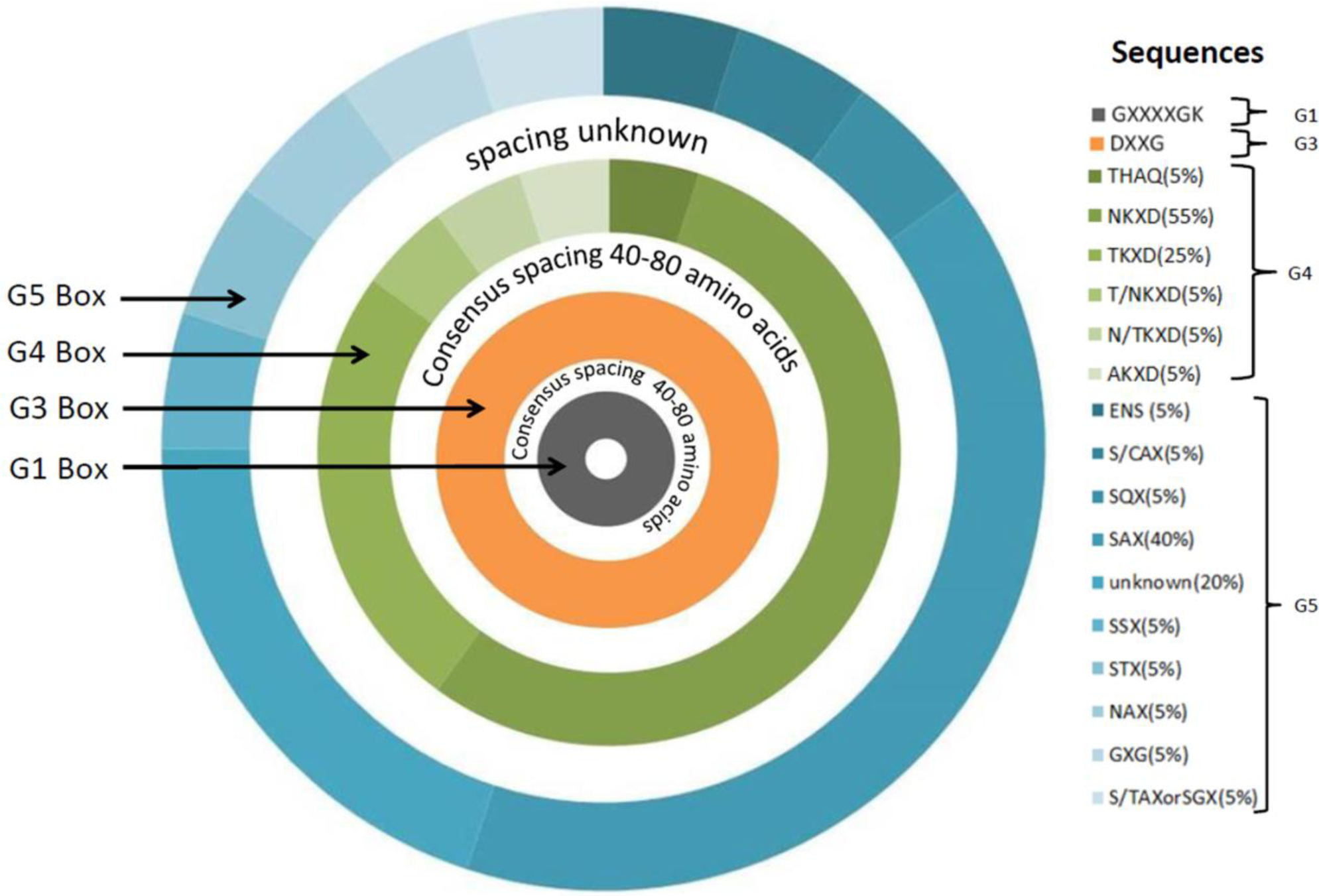
Doughnut plot representing consensus spacing between adjacent G boxes and consensus G box sequences across G protein families, collected by manual curation. Each concentric circle in the doughnut plot represents consensus sequences of different G boxes. G1 box (GXXXXGK, innermost grey circle), G3 box (DXXG, orange),G4 box (different shades of green), G5 box (outermost circle, different shades of blue).The white space in between consecutive concentric circles represents the amino acid spacers between consecutive boxes.

### Prediction of GTP-binding domains by Spacers and Mismatch Algorithm (SMA)

An algorithm, named “Spacers and Mismatch Algorithm” (SMA), was written (Figure 3) to predict G domains for the member proteins of the twenty different G protein families, using the manually curated G protein family-specific constraints for G domains (Suppl. Table 3).

**Figure 3:**
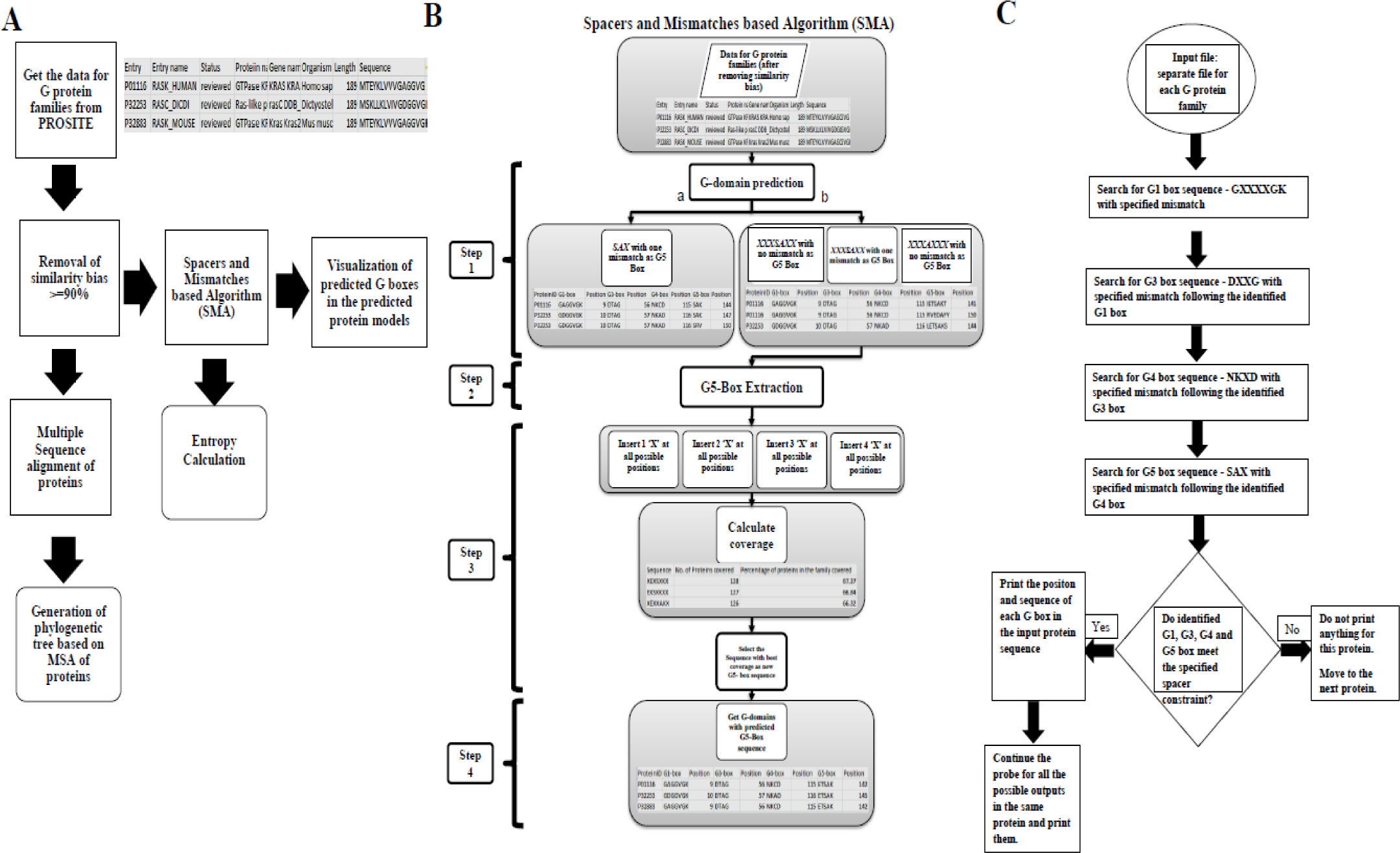
(A) Flowchart representing the entire methodology followed in the study (B) Spacers and Mismatch Algorithm (SMA) workflow (C) Details of G domain prediction by SMA (upto Step 1a)

SMA Step1a predicted G domains in more than 75% of the member proteins in each of the twenty G protein families [except for IRG (40%) and SRP (42.4%)] (Suppl. Table 5, columns 2 & 3). The rest of the proteins in each of the twenty G protein families probably did not give output due to differences in spacer length [e.g. in P24498 (Ras family), the spacer between G1 and G3 is 13 residues longer than the putative spacing (Suppl. Table 3)] or different length (e.g. Q91079 of the Ras family is only 93 residues long and does not appear to have a G1 or G4 box).

However, for many experimentally studied G proteins, multiple possible G domains were predicted for a single protein due to various possible combinations of predicted G boxes (G1, G3, G4 and G5). Figure 4 shows the individual G boxes, predicted using SMA, for two proteins each from Ras, Era and translational G protein families. These families were chosen to represent the findings because the G domains are very similar amongst homologs in the Ras protein family, (e.g., NRAS in humans and mouse), are very diverse in the translational family (e.g., EF2 in humans and mouse) while the Era family proteins have not been studied extensively.

**Figure 4:**
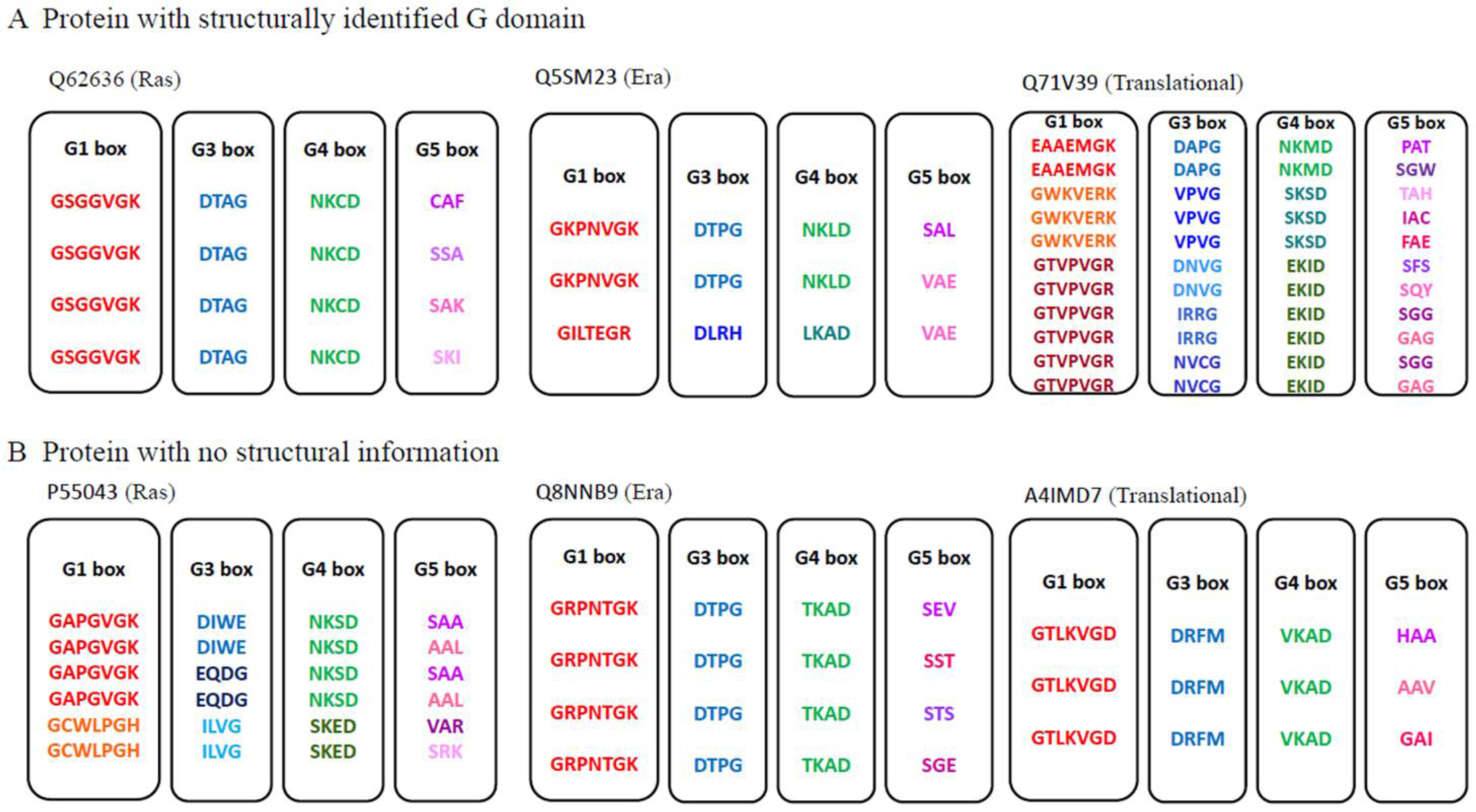
Prediction of multiple G domains per protein. (A) Q62636 (Ras), Q5SM23 (Era) and Q71V39 (translational) are proteins with structurally identified G domains (B) P55043(Ras), Q8NNB9 (Era) and A4IMD7 (translational) are proteins with no structural information. For each G box, different predicted motifs were depicted in different colors

One protein from each family [Q62636 (Ras), Q5SM23 (Era) and Q71V39 (translational)] has an experimentally identified G domain (PDB structures available) while the other protein [P55043(Ras), Q8NNB9 (Era) and A4IMD7 (translational)] has no available structural information (Figure 4). In the SMA generated output for proteins with solved structures, one of the three outputs for Q62636 and one of the four outputs for Q5SM23 matched exactly with the experimentally identified G domain. On the other hand, for Q71V39, the entire G domain was not predicted correctly in any of the 11 SMA outputs although 2 of the outputs was able to predict the G3, G4 and G5 boxes correctly.

Overall, it appears that SMA predicts multiple G domains either due to many predicted G5 boxes [e.g. Ras (Q62636); Era (Q8NNB9); Translational (A4IMD7) or many predicted G1-G5 boxes [e.g., translational (Q71V39), Ras (P55043), Era (Q5SM23)]. It was thus noted that most variability was observed in the prediction of the G5 box by SMA.

### SMA predicts significantly different G5 box motifs for some G protein families

“SAX” has been accepted as the G5 box consensus sequence (1, 3) since most experimentally discovered G5 boxes, including the exceptions to “SAX” across all G protein families, have been found to have either Ser or Ala interacting with the guanine ring of the GTP molecule. However, structural studies have identified many variations to the G5 box motifs across G protein families. For example, “SGL” (4ZKD), “TAL” (2H5E), “SSK” (4U5X), “GVG” (2OG2), “GEG” (1OKK), “SCA” (1E0S), “SSA” (1MOZ), “NAT” (2ZEJ), “SQL” (3W6N), “STR” (3HYR), “GVVNRSQ” (1JWY) etc.

It is to be noted that Step 1a of SMA, which searches for a G domain in a protein sequence using the G5 box constraint of “SAX” with one mismatch within the specified G4-G5 spacer length, essentially boils down to searching for only one amino acid i.e either S or A. This is because one mismatch will replace either S or A in one independent search. The variability observed in G5 box prediction by SMA (Figure 4), may be attributed to this obvious non-specificity in the search parameters. Thus, we decided to search for a more specific and probable family-specific consensus sequence for the G5 box.

To increase the precision to G5 box prediction, the search space was increased by increasing the length of the G5 box search string from 3 to 7 residues. SMA Step 1b (See Methods for details), searched for all possible 7mers after the predicted G4 box, that had either S or A in the centre and lay within the family-specified G4-G5 spacer range. Steps 2 and 3 of SMA were then performed to predict more precise G5 box consensus sequence per G protein family.

Interestingly, the output results indicated that there was no common 7-mer sequence that occurred in most members of all G protein families. Even closely related families like Ras, Ran, Roc and Arf showed different 7 mer sequences. On the other hand, each G protein family (except translational, TrmE and SRP) had one or more 7mers that occurred in >25% of the proteins of an individual G protein family. This indicated that there was probably no consensus G5 motif that was conserved across all G protein families, unlike the consensus motifs for G1 (GXXXXGK), G3 (DXXG) and G4 (NKXD) boxes.

An arbitrary threshold of 25% was chosen to ensure that the predicted 7mer covered at least a quarter of the members in a given G protein family. The motif which had the highest coverage was selected as the new consensus G5 box sequence for the given G protein family. Also, if there were Xs at the two ends of the predicted 7-mer, they were trimmed to give a more concise G5 box consensus sequence. Thus, the predicted family-specific G5 box sequences had different lengths. For example, EXSAX in the Ras family, SXXSXXS in the Roc family and ADXP in the OBG family (Suppl. Table 6).

### Comparison of G domains generated using “SAX” as consensus G5 box sequence versus using SMA-predicted G5 box sequence

#### (a) Prediction of fewer G domain boundaries per protein

SMA Step 4 (Figure 3b) was performed to search for all possible G-domains in each G protein sequence, using the family-specific constraints of consensus sequence, spacers and allowed mismatches listed in Suppl. Table 3 along with the additional constraint of using the new G5 box sequences predicted in Step 3.

The G1 box and the G5 box form the boundaries of the G domain in any G protein. The number of predicted G domain boundaries per G protein was significantly reduced (Figure 5) upon using the SMA-predicted G5 box sequence (from SMA Step 3; referred to as “SMA3” from now on), as compared to the output using “SAX” as G5 box sequence (from SMA Step 1a; referred to as “SMA1a” from now on). The Ras (members have similar G domains), translational (members have diverse G domains) and Era (less structural information) protein families were chosen to represent the findings. Figure 5 illustrates that for individual G proteins (vertical line) from all 3 G protein families, SMA Step 3 (Figure 5b) consistently reduced the number of predicted G5 boxes (red) and consequent C-terminal G domain boundaries while the number of predicted G1 boxes (blue) remained similar.

**Figure 5:**
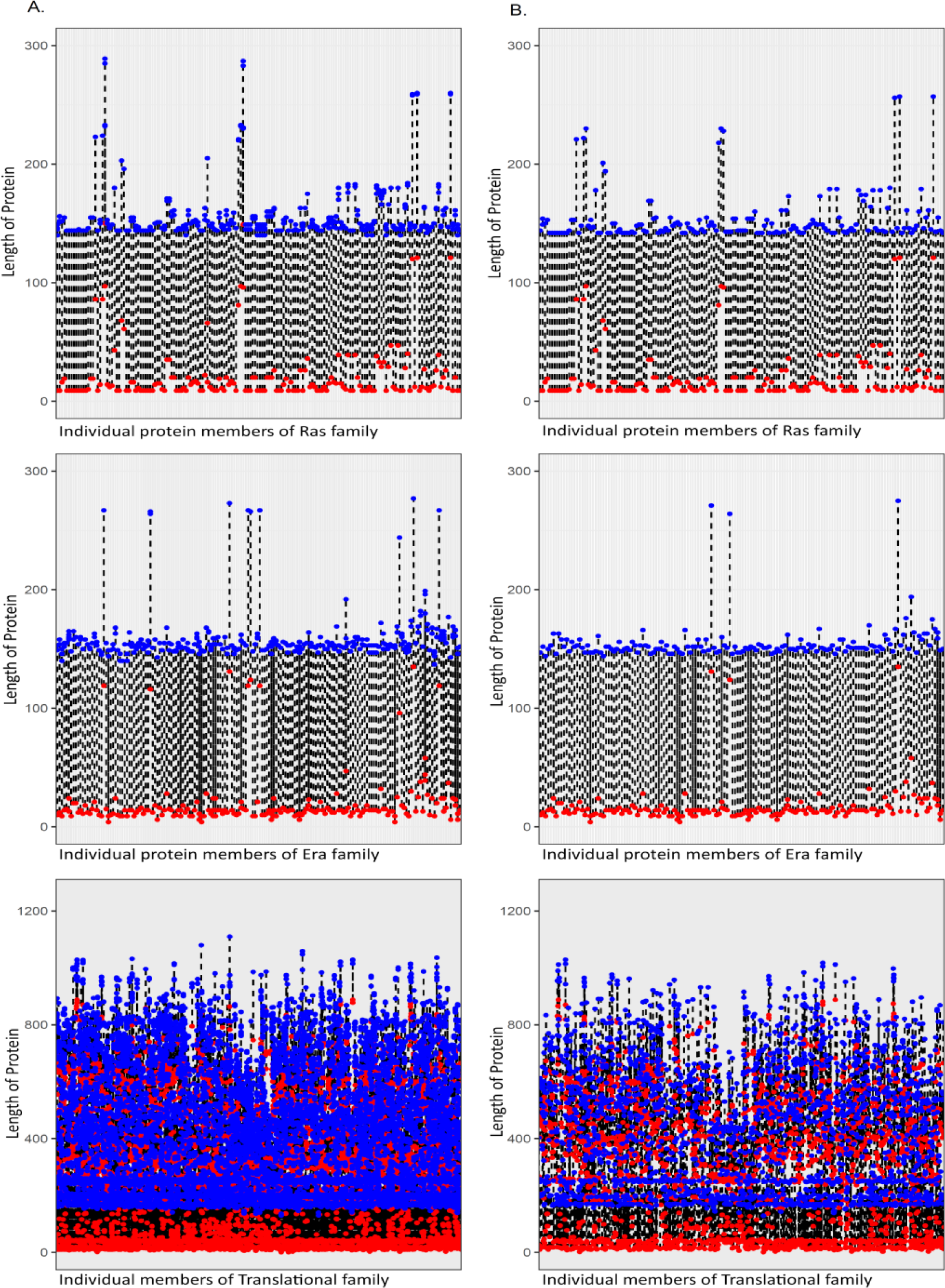
Comparison of predicted G domain boundaries (in Ras, Era and translational G protein families), before and after SMA-Step 3. Number of G domains predicted per protein (A) using “SAX” as G5 box (B) using newly identified G5 box after SMA-Step3. The X axis represents all the proteins in the G protein family. Each vertical line on the X axis represents each individual protein of the family. The Y axis represents the length of proteins (amino acids). Red dot marks the start of the predicted G1 box and blue dot marks the start of predicted G5 box. Each dotted line represents an individual G domain predicted per protein.

#### (b) Prediction of fewer unique G boxes per G protein family

We then investigated if SMA Step 4 had any effect on the prediction of all G boxes along with the prediction of fewer G domain boundaries and G5 boxes per G protein family. Upon using SMA3-predicted G5 box sequence (as compared to the output from SMA-1a), the predicted number of unique G boxes (G1, G3, G4 and G5 box) were significantly less i.e. >=50% in AIG-1, GB1/RHD3, Septin, IRG, Roc, EngB, FeoB, HflX, OBG, Rac, Arf and SRP families while no reduction was seen in the Ran family (Suppl. Table 7). Figure 6 illustrates the prediction of fewer unique G boxes in Ras, translational and Era G protein families.

**Figure 6:**
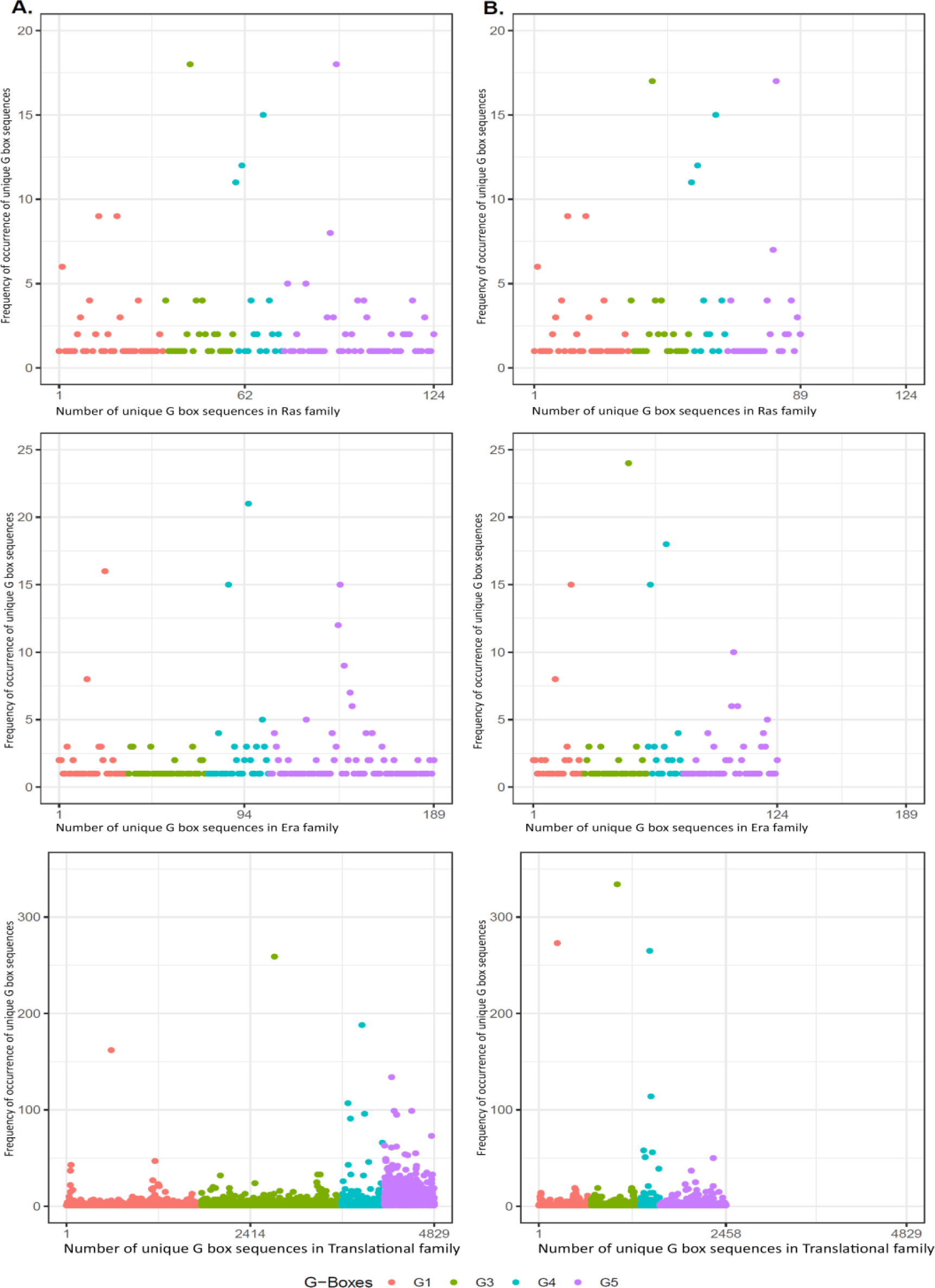
Comparison of unique G boxes (in Ras, Era and translational G protein families) predicted before and after SMA-Step 3. Number of unique G box sequences predicted (A) using “SAX” as G5 box (B) using newly identified G5 box after SMA-Step 3. X axis = total number of unique G boxes predicted in a G protein family; Y axis = frequency of occurrence i.e. the number of proteins (in G protein family), in which the predicted unique G box occurs). G1 (peach), G3 (green), G4(blue), G5(purple).

The prediction of fewer unique G boxes in AIG-1, GB1/RHD3, EngB, FeoB, HflX, Rac and Arf G protein families may be due to less proteins generating output upon using the SMA3-predicted G5 box sequence (Suppl. Table 5, columns 4 and 5). This again could be because in SMA Step3, the family specific G5 box was chosen based on its occurrence in >=25% member proteins of its family.

#### (c) Spacings between consecutive G boxes before and after SMA analysis

Spacers between consecutive G boxes were calculated in the G domains (predicted after SMA Step 3) of proteins in each G protein family. These spacers were compared to the spacers obtained after SMA Step 1a, in a representative histogram plot (Figure 7) for (A) Ras, (B) Era and (C) translational G protein families.

**Figure 7:**
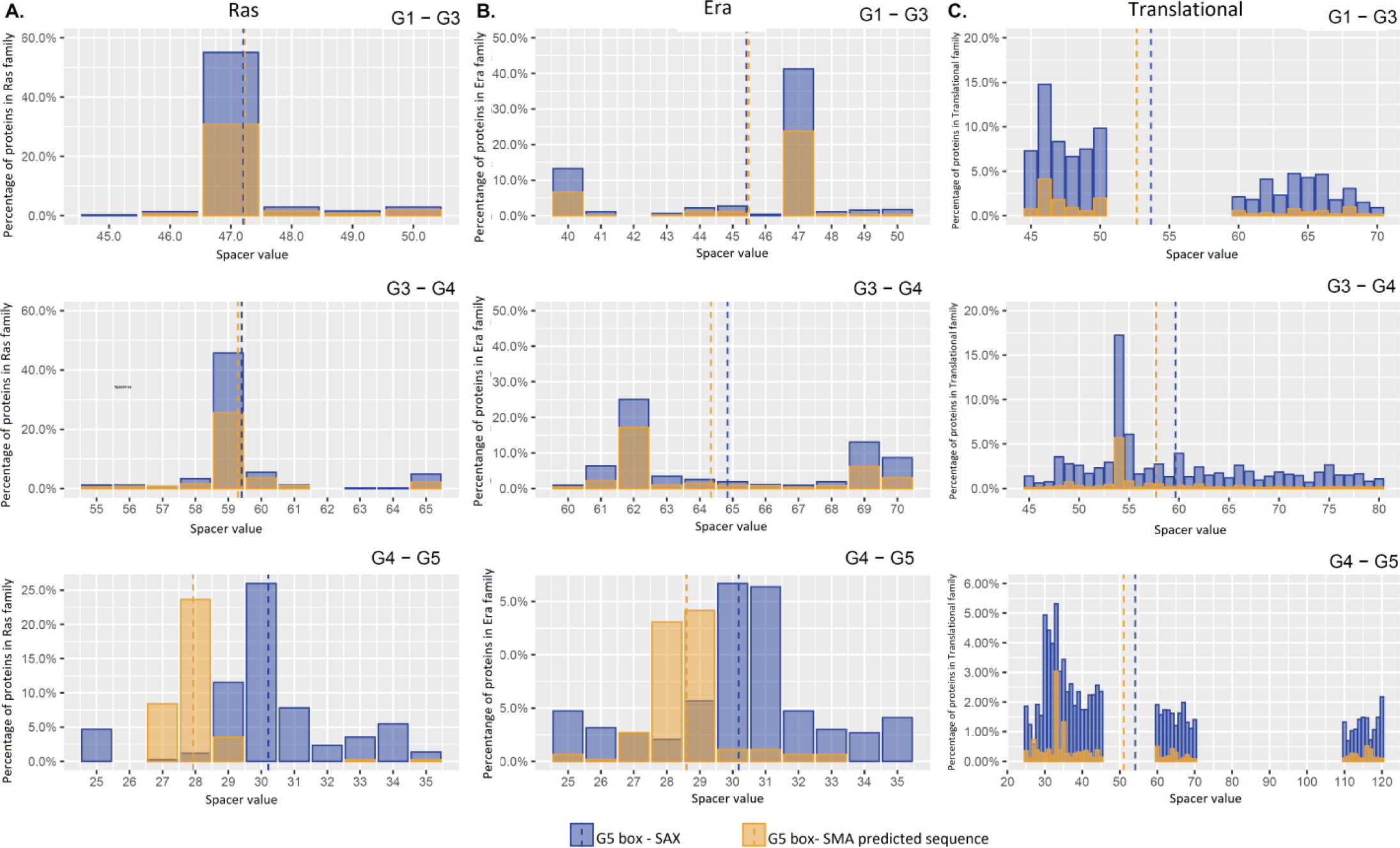
Comparison of amino acid spacers between consecutive G boxes predicted before and after SMA-Step 3. (A) Ras, (B) Era (C) translational families. X axis = range of spacer length (amino acids); percentage of proteins in G protein family, having indicated spacer value (bars); mean of inter-box spacing (dashed line) before (blue), and after (yellow) SMA-Step 3.

The mean of inter-box spacings was calculated by dividing the sum of individual spacer values (number of amino acids) in the defined spacer range by the total number of spacer values, where each spacer value represents the inter G box spacer in individual predicted G domains in a G protein family.

The percentage of proteins in G protein family, having indicated spacer value (bars); mean of inter-box spacing (dashed line), before (blue), and after (yellow) SMA-Step 3, varies between families as well as between specific spacers. These parameters are either identical in Ras (G1-G3, G3-G4) and Era (G1-G3) families (Figure 7A, 7B), quite similar in Era (G3-G4) and translational (G1-G3, G3-G4 and G4-G5) (Figures 7B and 7C) or very different in Ras (G4-G5) and Era (G4-G5).

The percentage of proteins having similar spacers before and after SMA-Step 3 were found to be similar in all the G protein families except for Hflx, FeoB and SRP (for G1-G3 spacer), FeoB and SRP (for G3-G4 spacer) and AIG1, Arf, EngB, Era, IRG, HflX, OBG, Rab, Rac, Ran, Ras, Septin, SRP, TrmE (for G4-G5 spacer) (Suppl. Table 8).

Thus, it appears that in a large number of G protein families, the percentage of proteins having similar G4-G5 spacers as well as the mean of the G4-G5 predicted spacings, differs significantly before and after SMA3. However, the G1-G3 as well as the G3-G4 spacers do not differ in most families. This implies that there is a significant difference between the SMA3-predicted G5 motif and “SAX”, which was the accepted G5 consensus.

#### (d) Multiple Sequence alignment based phylogenetic relatedness of predicted G domains

Phylogenetic tree analysis constructed using full length protein sequences of individual G protein family members (Figure 1) illustrated that G proteins that belong to the same family may not always cluster together. This may be because most of the G proteins, except the Ras superfamily members, are characterised by multi-domain proteins (one of these domains being the conserved G domain). To investigate this further, phylogenetic trees were constructed (Suppl. Figure 4) for just the predicted G domains (before or after SMA Step 3) of the same five representative proteins per G protein family (which were studied in Figure 2).

Overall, G domains (predicted after SMA Step 4) of G proteins that belong to the same family tend to cluster together more strongly (Suppl. Figure 4b) as compared to G domains (predicted after SMA Step 1a; Suppl. Figure 4a) or full length G proteins (Figure 2). This is especially true for members of Roc, AIG1, translational and IRG G protein families. For these families, the differences observed between Figure 2 and Suppl. Figure 4 may be attributed to the insertions within the G domain in some translational and AIG-1 proteins as well as additional domains in the full-length G proteins.

### Prediction of G protein family specific G1, G3 and G4 box consensus sequences

SMA Step 3 predicted G protein family-specific G5 box consensus sequences, that were previously not reported (Suppl. Table 6). It was intriguing to then ask if there were also G protein family-specific G1, G3 and G4 box motifs beyond the consensus”GXXXXGK”, “DXXG”,”NKXD sequences respectively. In other words, can the consensus motifs be modified and more specifically, are there any family-specific patterns, which could possibly give better representation of the “X” residues (where X=any of the twenty amino acids).

G1, G3 and G4 box sequences, were predicted for all the proteins in each G protein family after SMA Step 4 and visualized as sequence logos. More distinct G protein family specific G box consensus motifs were predicted in most G protein families except AIG-1, GB1/RHD3, OBG, Roc and translational families. G protein family specific G box consensus motifs of all the proteins in Ras, Era and translational protein families are shown in Figure 8. For example, the first “X” in G3 box is found to be “T” in all three protein families whereas, the second “X” in G3 box can either be “P” or “A” (Figure 8). Several interesting family-specific signatures were observed. For instance, the predicted G3 consensus was “DXXS/D” in the AIG1 protein family but “D/WXXG” in Arf while the predicted G4 consensus was “TKXD”in Dynamin, SRP, Rac and EngB G protein families and “AKXD” in Septin (correlates with corresponding PDB structures). Some families had very specific predicted G box consensus motifs. For example, G3 box (DLPG) in FeoB, G1 (GESGAGK) and G4 (TKVD) boxes in IRG and all four G boxes (GDGGTGK, DTAG, NKVD and SAKSN) in Ran. (Suppl. Figure 5)

**Figure 8:**
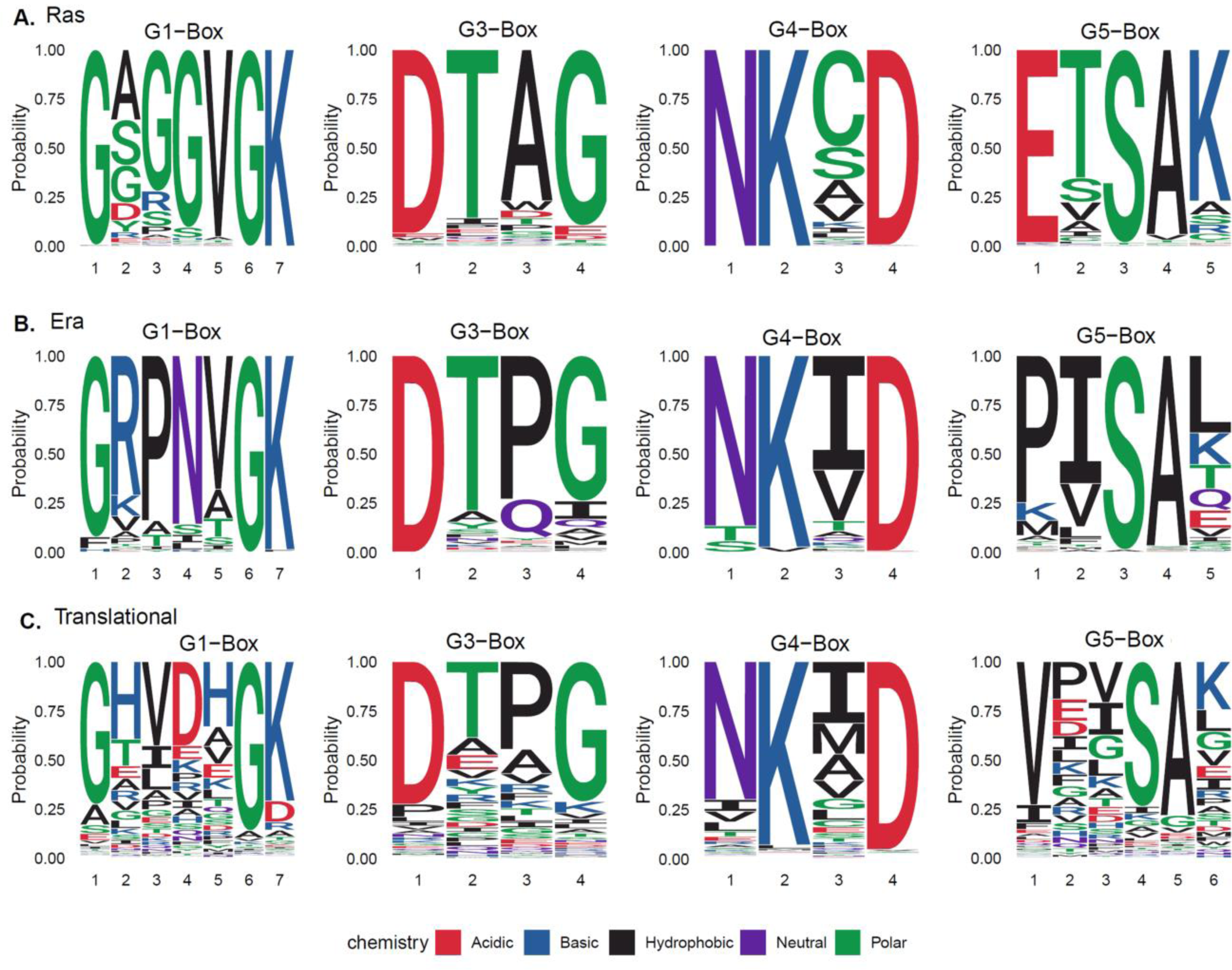
G protein family-specific G box motifs predicted using SMA3. (A) Ras, (B) Era (C) translational families. X axis= amino acid positions in respective G box; Y axis= probability of occurrence of an amino acid from 0 to 1.

The diversity of predicted G boxes in the translational family is consistent with the documented sequence diversity of its member proteins; similar G box diversity is also seen in Roc, OBG, AIG-1 and GB1 families (Suppl Table 1, Figure 1, Suppl. Figure 4). On the other hand, the similarity of predicted G boxes in the less studied Era family members, may imply greater homology in their G domains since Ras family members, which have high homology, also appear to have very similar predicted G boxes.

It is interesting to note that the amino acids at the amino and carboxy-terminal ends of each G box were conserved across all families; these conserved residues were in fact, parts of the previously accepted G box consensus motifs which had defined amino acid signatures instead of “X”. [i.e, the G and GK “GXXXXGK”(G1), D and G in”DXXG” (G3) and NK and D”NKXD (G4)].

In order to test the specificity of the SMA output, it was validated by using proteins, other than canonical GTP binding proteins, as input (Suppl. File 4). For example, no output (after SMA Step 1a) was generated when hemoglobin chain A or hemoglobin chain B were used as query sequence. Moreover, ATP binding proteins (Eg; RBBA protein in *E.coli*) which have a P loop (G1 box) but no other G box, do not generate any output. In proteins like human OLA1 which bind to both ATP and GTP, the generated output has one combination of predicted G boxes (G domain) that binds GTP.

To maximise the number of predicted G domains per G protein, the family-specific G box consensus sequence and inter G box spacer constraints used to run SMA were kept broad. In order to test the significance of the SMA results and to ensure that the broad constraints did not end up predicting G domains in any string, a negative control experiment was performed. Suppl. Fig. 6 illustrates that after shuffling the protein sequences, the number of G-Domains predicted per sequence (red) becomes significantly less. This further validates the specificity of SMA prediction.

### The predicted G box sequences does not depend on the size of the G protein family

Figure 8 demonstrates that for certain G boxes in some G protein families, there is significant diversity in individual positions within each box (e.g. translational family). It can be argued that this observed diversity may be due to the varying sizes of the different G protein families i.e a protein family with more members would show more variation in G box motifs merely because of larger representation than a protein family with fewer members. To probe whether the diversity of predicted G box sequences was a function of the size (number of member proteins) of a G protein family, the entropy of individual G boxes was calculated for each family. The calculated entropy values gave insights about how random the “X” position in each of the G boxes (i.e. GXXXXGK” for G1, “DXXG” for G3 and “NKXD” for G4) is and how randomness of individual “X” positions impact the total entropy of the G-Box.

Figure 9 investigates the correlation between the total entropy of each G box and the size of each G protein family. The Translational family is the largest G protein family with 2870 member proteins; however, the entropy calculated for each of its G boxes is not significantly high. On the other hand, small G protein families like Roc, with only 20 members, demonstrate highest entropy for G1, G3 and G5 boxes while the GB1/RHD3 family, with 50 members, has the highest entropy for G4 box. The IRG (11 members), FeoB (19 members) and Ran (30 members) families have the lowest entropy for G1, G4 and G3, G5 boxes respectively in spite of having similar sizes to that of Roc and IRG families (Figure 9). Thus, it appears that the observed variation in G box motifs does not correlate with the size of the protein family but maybe because of other properties of individual proteins.

**Figure 9.**
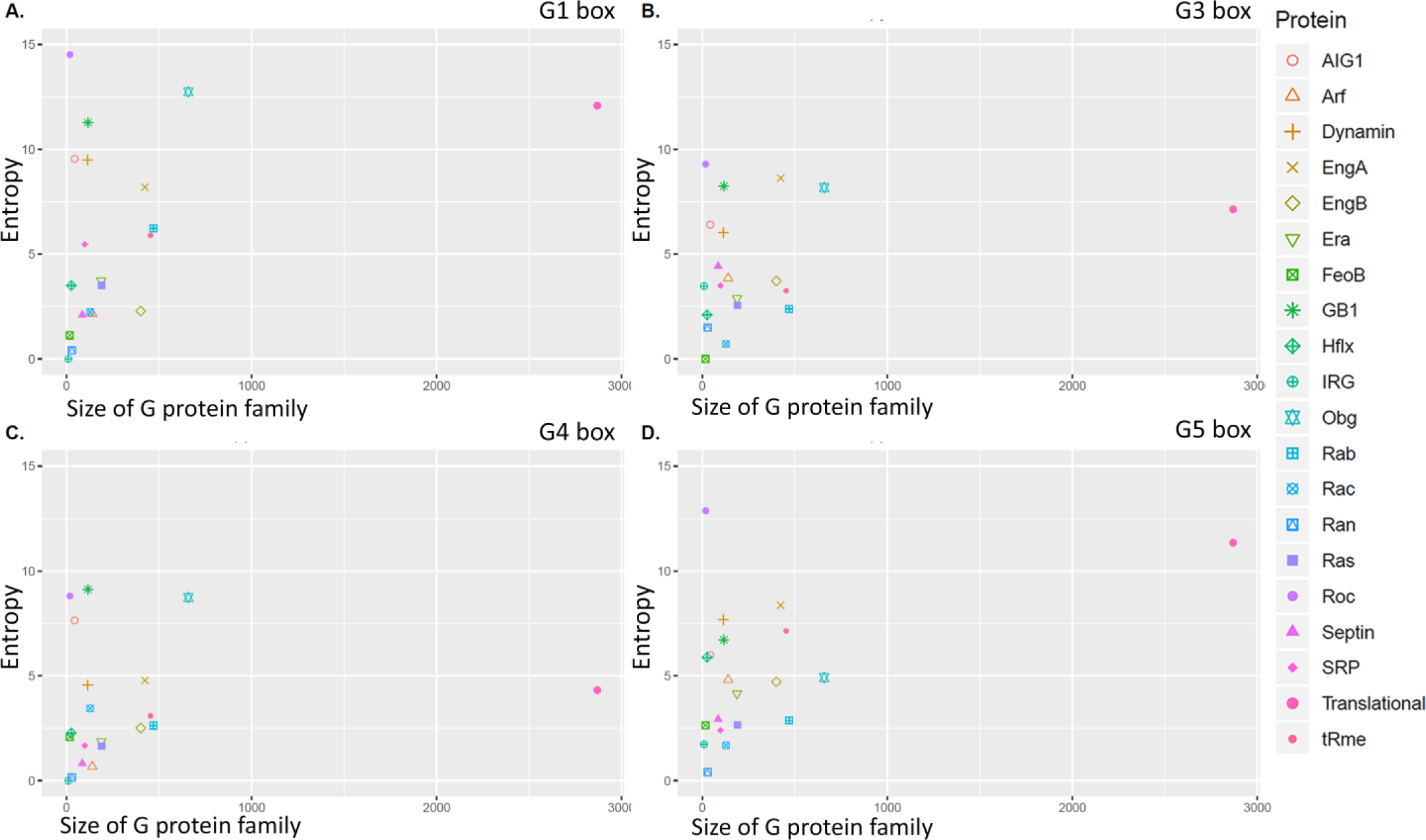
Correlation between G box entropy and G protein family size: Scatter plots where the X axis represents the total number of proteins (size) per G protein family and the Y axis represents total entropy of each G box (A) G1 box (GXXXXGK), (B) G3 box (DXXG), (C) G4 box (NKXD) (D) G5 box predicted by SMA. Each G protein family is represented by a different color as shown in the key.

### Position of predicted G5 box sequence on the loops of available structures or I TASSER predicted structures provide confidence to the sequence based G5 box analysis

G boxes are the most conserved regions of G domains (3). The conserved G boxes of G proteins occur in loop regions (8), which are the least ordered secondary structural elements of any protein structure (36). This is unlike transcription factors, in which most conserved sequences fall on alpha helices or the lipocalin protein family (6) which have conserved beta sheets. We decided to test whether the sequence-based prediction of G protein family-specific G box sequences mapped on to loop like structures of the corresponding protein.

Predicted G box motifs (using the newly identified G5 box sequence) were mapped onto available protein structures downloaded from the PDB database [Fig.10a; Q62636 (Ras), Q5SM23 (Era) and Q71V39 (translational)] as well as I TASSER predicted protein structures [Fig.10b; P55043(Ras), Q8NNB9 (Era) and A4IMD7 (translational)] from Ras, Era and translational G protein families. Figure 10 demonstrates that the predicted G box sequences mapped to the loop regions in both the available (Fig.10a) and predicted (Fig.10b) protein structures. This structural alignment provides confidence to the sequence-based G box prediction. Additionally, the fold of the G domain is found to be very similar between the two proteins of the same G protein family [Q62636 and P55043(Ras), Q5SM23 and Q8NNB9 (Era) and Q71V39 and A4IMD7 (translational)].

**Figure 10:**
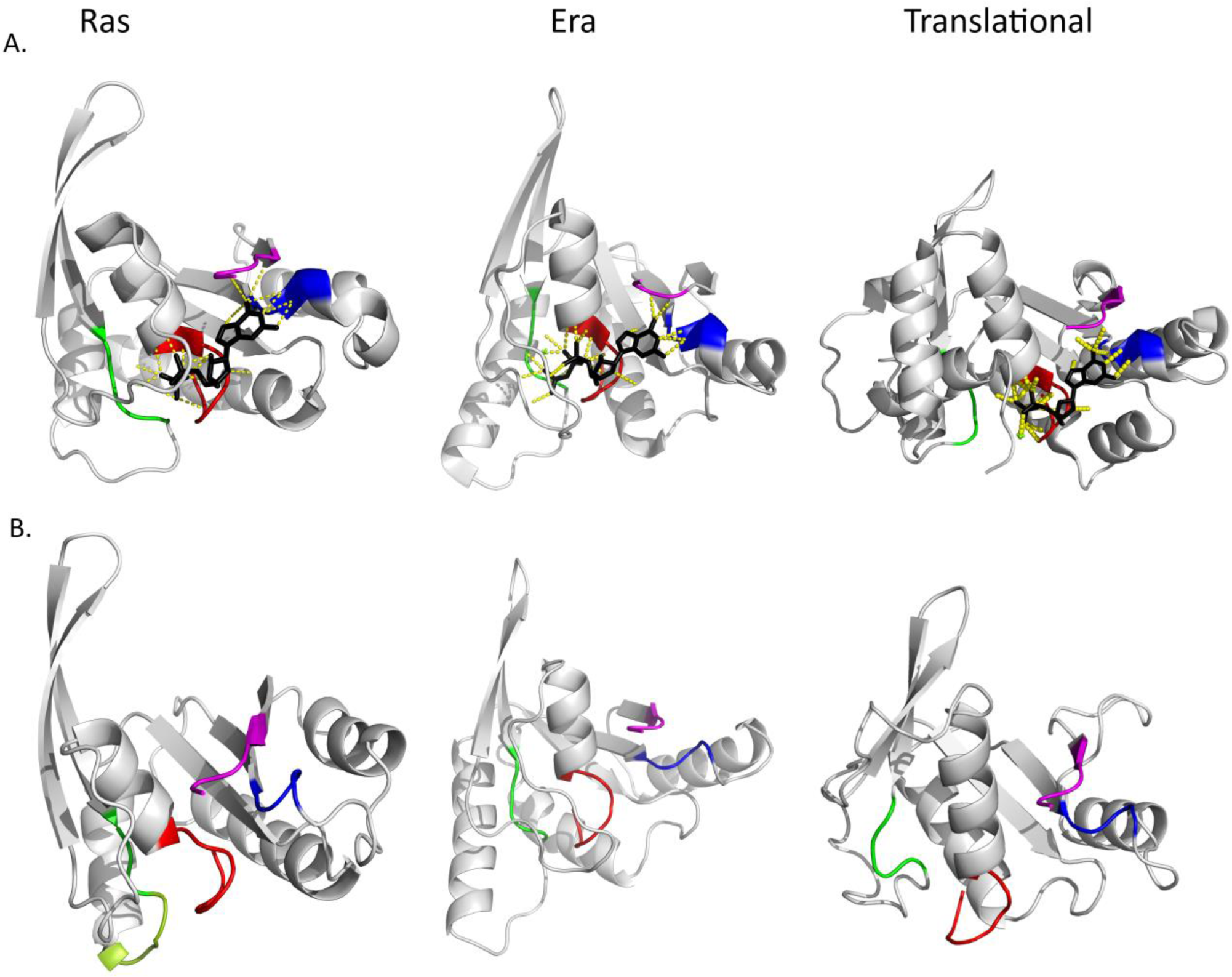
Superposition of the predicted G boxes to the secondary structure of G proteins. Legend: Predicted G domains (gray), visualized using PyMol, were mapped onto (a) available protein structures downloaded from PDB [Q62636 (Ras), Q5SM23 (Era), Q71V39 (translational)] (b) I TASSER predicted protein structures [P55043(Ras), Q8NNB9 (Era), A4IMD7 (translational)]. G1 box (red), G3 box (green), G4 box(blue), predicted G5 boxes (magenta). GTP/GDP are shown in black in (a).

## Discussion

The GTP binding domains of all G proteins have been characterized by a set of consensus G box sequences (G1-G5) and inter-box spacers (3, 7). However, with the recent identification of many G protein families, it is observed that there is a vast diversity in the sequence of G proteins. This diversity is reflected in the sequences of the G-boxes and more specifically in inter-G box spacer length and the G5 box consensus sequence. Here, we come up with a tool called SMA, which can identify G protein family-specific G box sequences and inter-G box spacer length, based on the constraints within individual families.

SMA is a one-of-a-kind algorithm (Figure 3a, b,c), which can perform an approximate search for multiple sequences occurring with a given range of gap between them. In other words, unlike tools like GTP binder (30), ELM (29), ScanProsite (31), Pfam (32) etc., SMA offers the advantage of using multiple proteins in the input file and searches for all G boxes simultaneously instead of one after the other. It uses consensus patterns with allowed variations and maximum and minimum gaps allowed between each pattern. Moreover, all the constraints in the algorithm are customizable and the user has the flexibility to change the G box sequence including the number of allowed mismatches as well as the inter G box spacer length. This is especially useful, if the protein of interest has very low amino acid sequence similarity with well-studied GTP binding proteins and/or is less studied i.e. little or no information is available from structural or biochemical studies.It is to be noted that SMA is a generalized algorithm, which is not restricted to finding amino acid motifs in protein sequences. The algorithm can be appropriately modified to identify patterns in any sequence including those of biological importance like DNA or RNA.

An exhaustive analysis of twenty different PROSITE G protein families, demonstrated that G box sequences and inter-G box gaps are often class-specific. Thus, it is proposed that G protein families could possibly be classified based on family-specific G box sequences and inter-G box spacer lengths. Moreover, SMA was able to come up with better representations of the family-specific G box motifs. For instance, it was possible to predict if the “X”s in the established G1 box motif (GXXXXGKS/T) could be better represented by a specific set of residues rather than any of the twenty residues (Figure 9). SMA, was also able to predict G5 boxes for the different G protein families. This was significant since the G5 box marks the carboxy-terminal boundary of a G domain and hence demonstrates that SMA can be used to predict more precise boundaries of a G domain, within a large multi-domain protein.

Analysis of the sequence divergence between different G protein families may provide mechanistic insights into the differential basis of GTP binding and hydrolysis in different G proteins like Ras and Dynamin. For instance, Dynamin proteins have low nucleotide affinity and low GTP hydrolysis rates with a GTPase cycle that is regulated by nucleotide binding and subsequent dimerization (GAD) as compared to Ras proteins which have low intrinsic nucleotide exchange rate, high nucleotide affinity and are regulated by GEF, GAP and GDI proteins (21). It is tempting to speculate if these differences can be attributed to the difference in inter G box spacers (Suppl. Table 3), G4 motifs (Ras; “NKXD” in 121P,1×1R and Dynamin; “TKXD” in 1DYN, 1JWY) and G5 motifs (Ras; “SAK” in 121P,1×1R and Dynamin; “SQK” in 1DYN, “GVVNRSQ” in 1JWY) between these two families. Interestingly, fusion Dynamins like Mfn1/2 lack the consensus G5 box while the consensus G5 box is “SQX” in fission Dynamins like Drp1 and human Dynamin -1.

Some G proteins have the ability to bind and/or hydrolyse more than one nucleotide apart from GTP. This may be explained by the presence of additional domains (for example, the Hflx proteins, which have a G domain as well as an ATP binding ND1 domain (22). Alternately the nucleotide specificity may be altered by changes in the G4 box motif (“NKXD”). For example, within the OBG family, the G4 motif is “NLXD” in human YchF/OLA1, which functions as an ATPase rather than a GTPase (37) and “NhXE”(where h is a hydrophobic residue) in *Oryza Sativa* YchF, which binds both ATP and GTP with similar binding specificities (38).

The EngA/Der/YfgK/YphC family is characterized by two tandemly occurring GTPase domains separated by an acidic linker (20). SMA’s ability to independently predict two distinct domains along with corresponding G boxes in individual EngA members, provides confidence to the method. Moreover, the precision of SMA prediction was further validated by the fact that no G domain was predicted in the center of the EngA proteins.

It is also interesting to note that some G proteins have different insertions within the G domain For example, eIF-2***γ*** has a zinc ribbon insertion between the G2 and G3 boxes (3) while AIG-1 proteins have a hydrophobic motif insertion between G3 and G4 boxes (13). SMA predicts structurally identified G boxes and the corresponding G domain for some of the AIG-1 and eIF2 proteins. SMA predictions for these families may improve as more structural information becomes available and the SMA parameters can be changed accordingly.

Detailed analysis of SMA results can also be used to explore interesting evolutionary relationships amongst homologs of the same protein. For example, Rho1 protein in *E. histolytica, B. vulgaris, P. sativum* does not give any output with specified G box sequences and inter G box spacers. This is because, in *E. histolytica* Rho1, the putative G5 box (EASSV) is only 28 amino acids away from the putative G4 box (LKVD) which differs from the defined G4-G5 spacer range of 40-50 amino acids, that is present in most Rho1 homologs (suppl. Table 3). Rho1 from *B. vulgaris* and *P. sativum*, lacks a putative G3 (NKXD) box and the putative G5 box (ECSSK) is 37 amino acids away from the putative G4 box (TKLD) which differs from the defined G4-G5 spacer range (40-50 amino acids). On the other hand, SMA predicts identical G1 (GDGACGK), G3 (DTAG) and G5 (ECSAK) boxes in Rho1 homologs from *D. melanogaster, A. gossypii, C. elegans, S. pombe, K. lactis, S. cerevisiae* and *C. albicans* but different G4 box sequence (NKXD in *D. melanogaster* and *C. elegans* and CKXD in *A. gossypii, S. pombe, K. lactis, S. cerevisiae* and *C. albicans*). These observations could be further investigated to understand the nucleotide binding properties of the different Rho1 homologs.

It was also noted that for certain G proteins, the inter G box spacer length was conserved within homologs present in the same kingdom of life but differed between kingdoms. For example, the IF2/eIF5B proteins of the translational family, which is functionally conserved in all domains of life, has respective G1-G3, G3-G4 and G4-G5 spacer lengths of 46, 54 and 34 or 36 in bacteria (Eg. IF2 structures from *B. stearothermophilus and T.thermophilus*), 67 or 72 or 77, 56 and 33 or 35 in Archaea (IF2 structures from *P. furiosus, P. abyssi* and *S. solfatricus*) and 64 or 72, 54 and 66 or 68 in Eukarya (Eg. eIF5B from *S. cerevisiae, M. thermautotrophicus*). Similar paralog-specific spacers are also observed in eukaryotic RFs (eRF3, EF1a2, ET-Tu) as well as the OBG family (OBG, DRG, NOG1, Ygr210 compared to YchF/YyaF). These interesting observations about diversity of inter G box spacers within the translational family, agrees with reported evidence of the divergence of translational factor subfamilies (39). This also suggests that certain families like the translational factors may be further classified into distinct subfamilies based on sequence divergence and possibly different cellular functions.

Our current observations are however, biased by the PROSITE classification of G protein families, our manual curation of family-specific G box sequences and our imposed constraints for allowed mismatches in individual G box sequences and inter G box spacer length. Thus, any exceptions to the above constraints will be missed and may need to be changed, as more structural information is obtained about members of different G protein families. Moreover, it is to be noted that SMA had an inherent bias in G5 box prediction since it only looked for patterns (with allowed mismatches) which had “SAX” in the centre. Also, the G5 boxes that are identified using a 25% threshold, miss under-represented putative G5 box sequences. This is specifically important for less-studied families like AIG-1, Era, GB1/RHD3, IRG and Roc, in which there is very little available structural information for individual members. On the other hand, for families like Ras, for which there is a wealth of structures, our imposed constraints may stand the test of time.

## Methods

### Selection of G protein families for further analysis

PROSITE is a collection of protein domains and families along with the structural or functional patterns associated with them. “Guanine nucleotide-binding” was used as the search string to find PROSITE [Pubmed id 23161676] entries corresponding to G proteins. 21 documentation entries were identified to have “Guanine nucleotide binding” profiles. 4 out of these 21 entries include proteins which interact with GTP binding proteins but do not bind to GTP themselves [PDOC00210 G-protein coupled receptors family 1, PDOC01002 G-protein gamma subunit, PDOC00574 Trp-Asp (WD-40) repeats and PDOC51717 Very large inducible GTPase (VLIG)]. <u>PDOC00017</u> ATP/GTP-binding site motif A (P-loop) includes proteins containing only the P loop and not the entire G domain. PDOC00962 Hydrogenases expression/synthesis hypA family does not have any GTP binding proteins. Also, PDOC51721 Circularly permuted (CP)-type proteins are excluded as this family consists of proteins that contain G boxes in a different order from the canonical G1-G2-G3-G4-G5 pattern, e.g. DAR GTPase 3 in *Arabidopsis thaliana* has G boxes in the order G4-G1-G2-G3 (40).

The Uniprot/Swiss-prot true positive sequences for each of the remaining 14 entries were selected for further analyses. These include the following guanine nucleotide-binding (G) domain signatures and profiles: PDOC51720 AIG1, PDOC51714 Bms1, PDOC00362 Dynamin, PDOC51712 EngA, PDOC51706 EngB, PDOC51713 Era, PDOC51711 FeoB, PDOC51715 GB1/RHD3, PDOC51705 HflX, PDOC51716 IRG, PDOC51710 OBG, PDOC51719 Septin, PDOC00273 Translational (tr), PDOC51709 TrmE. In addition to the above entries, the following entries, which were not represented in the results of search string “Guanine nucleotide binding”, were included: PDOC51417 (small GTPase family which includes all the canonical small G proteins like Arf, Miro, Ran, Rab, Ras, Rho/Rac, Roc and SAR1) and <u>PDOC00272</u> (SRP54 family, a SIMIBI class protein). Bms1, Miro and SAR1 families were excluded from this study due to the unavailability of sufficient structural information of any of their protein members. Also, since PROSITE does not have a separate documentation entry for alpha subunits of heterotrimeric G proteins, one protein from each of the four families of alpha subunit (namely Gi, Gs, Gq, G12/13) of heterotrimeric G proteins was chosen as a representative (Suppl. Table 9).

An excel file (with uniprot id, protein name, length of the protein, protein sequence and PDB structure information) was retrieved for true positive sequences of all the 20 PROSITE documentation entries from PROSITE and used as the input file for our analysis (Suppl..File 1)

### Removal of similarity bias from each PROSITE documentation entry

Levenshtein distance (edit distance) was used to compare all the member protein sequences of the same G protein family amongst each other. Only one sequence from all the sequences that showed >=90% similarity was retained to reduce redundancy and remove the similarity bias from further analysis. (https://github.com/RichaRashmi-projects/G-Protein-Project.git)

### Multiple sequence alignment of proteins and phylogenetic tree generation for all G protein families

After removal of similarity bias, FASTA files (containing the non-redundant protein members of each of the G protein families) were used to perform multiple sequence alignments followed by phylogenetic analysis. These were performed using ClustalW and maximum likelihood method respectively, provided in the MEGA X software package (41). The phylogenetic tree was visualized using iTOL (42).

### Spacers and Mismatch algorithm (SMA)

The Spacers and Mismatch Algorithm (SMA), uses a data-based approach to predict G domains for the GTP-binding proteins (Figure 3b, c). It takes special consideration to define a more precise G5 box and consequently a better carboxy-terminal boundary of a G domain. The detailed explanation of the workflow of SMA can be found in the Supplementary information.

### Significance testing of predicted G-domains

In order to test the significance of the SMA results, a negative control dataset was created by shuffling (fifty times) the protein sequences for each G protein family. The number of G-Domains predicted per protein (by SMA) was counted after each shuffle. After 50 shuffles, the total number of predicted G domains per sequence was averaged and compared with the total number of predicted G domains (after running SMA-3) in the actual protein sequence.

### Generation of protein models and visualization of predicted G boxes

Protein structures of one protein each from Ras (P55043), Era (Q8NNB9) and translational (A4IMD7) G protein families were predicted using I TASSER (44). Briefly, the amino acid sequence of these proteins were submitted to the I TASSER server with no specified template. I TASSER server identified suitable templates depending on sequence similarity searches from the PDB database. Monte Carlo simulations were used to assemble the full-length conformations of identified templates and models were generated for the sequence of interest. All the conformations were confirmed and cluster centroids were identified, which were then used to build the final models after refinement of cluster centroids.

In addition, one protein each from Ras (Q62636), Era (Q5SM23) and translational (Q71V39) G protein families were extracted from PDB.

The predicted G boxes for all six of the mentioned proteins were visualized as loops using PyMol (45, 46).

### Entropy Calculations

To investigate if the classes with more number of proteins have more variation in the individual G boxes, entropy of each G box was calculated using the entropy function in scipy (stats.entropy). This was performed by adding the entropy of each position of the predicted G box sequence. Scatter plots were generated where the X axis represents the total number of proteins (size) per G protein family and the Y axis represents total entropy of each G box.

## Acknowledgements

We acknowledge Dr. Umashankar Singh and Dr. Sairam S. Mallajosyula for providing valuable suggestions about the study. Special acknowledgements to Dr. Murali Krishna Enduri for helping HS understand and troubleshoot Fig. 3C, Dr. Althaf Shaik and Ms. Deepshikha Ghosh for their timely help with PyMol, Mr. Divyesh Patel for suggestions about trying MEGA for MSA and phylogenetic tree analysis.

## Figures

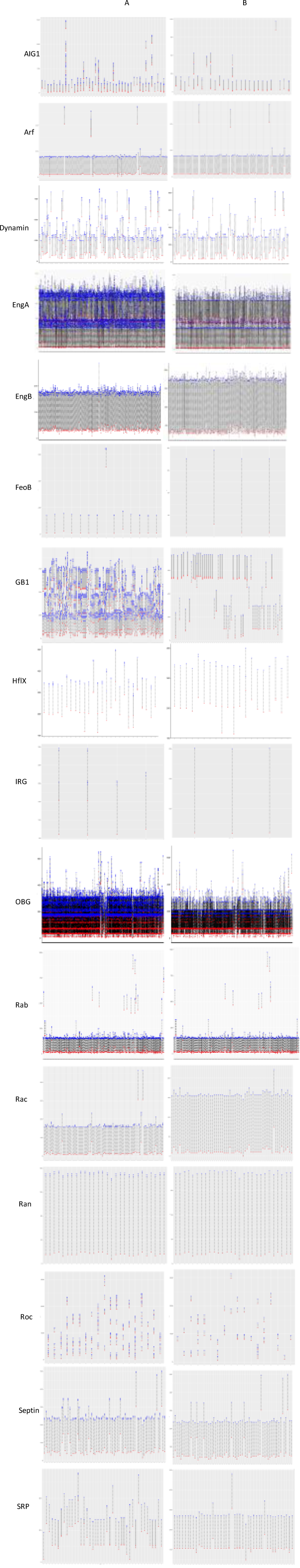

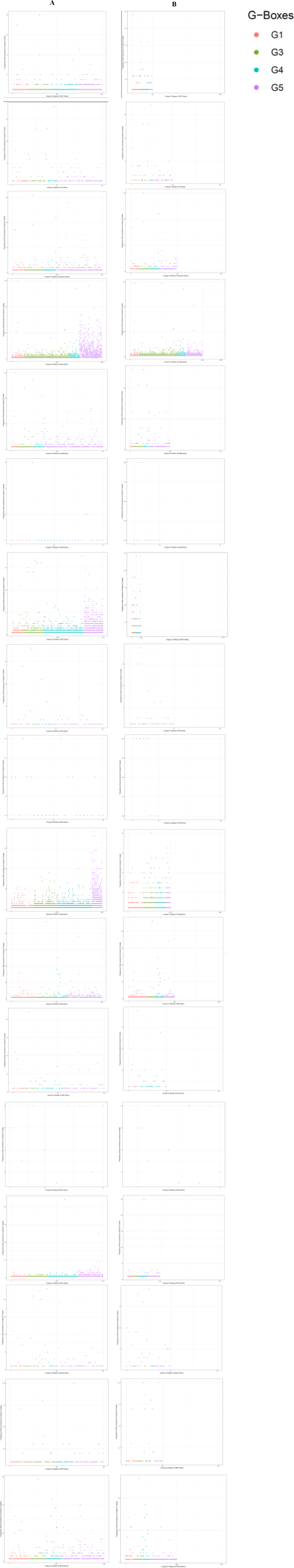

