## Supplemental methods, figures, tables for "G-domain prediction across the diversity of G protein families"

Hiral M. Sanghavi[1], Richa Rashmi[1], Anirban Dasgupta[2] and Sharmistha Majumdar1*
1 Discipline of Biological Engineering, Indian Institute of Technology Gandhinagar
2 Discipline of Computer Science and Engineering, Indian Institute of Technology Gandhinagar and Sharmistha Majumdar[1]*

*Sharmistha Majumdar

**This PDF file includes:**

Supplementary text
Figures S1 to S6
Tables S1 to S9

**Supplementary Information Text**

**Supplementary Methods**

### Spacers and Mismatch algorithm (SMA)

Step1- G-domain prediction

A python script (https://github.com/RichaRashmi-projects/G-Protein-Project.git ) was written, to search for all possible G-domains (along with putative G boxes) in each G protein sequence, using the family-specific constraints of consensus sequence, spacers and allowed mismatches listed in Suppl. Table 3. Each protein sequence was used as a string. Four different G boxes (sub-strings; 1.”GXXXXGK”, 2. “DXXG”, 3.”NKXD” and 4.”SAX”) were searched in the string, one at a time. After the identification of the first conserved amino acid of a G box, a sliding window was used to find the complete G box sequence. After all four G boxes (G1-G3-G4-G5) were predicted, the code checked to see if the defined spacings between the consecutive G boxes was maintained. This led to the prediction of the entire G domain with predicted G box sequences that had defined inter-box spacings. The sequences and start positions of each predicted G box were saved along with their respective protein IDs. The algorithm implemented in the script is shown in Figure 3.

Step 1 was implemented twice. (a), SAX with one allowed mismatch was used as the G5 box consensus sequence (Figure 3b) (b) Next, to predict a better and possibly longer G5 consensus sequence centred around SAX, we modified the search in 3 ways- (i) “XXXSAXX” with no mismatch (ii) “XXXSAXX” with one mismatch and (iii) “XXXAXXX” with no mismatch , where X can be any of the 20 amino acids (Figure 3c). Thus, G-domain identification for each of the 20 G-protein families, was performed again using these modified G5 box sequences.

Step 2- G5 box Extraction

Only the G5 results obtained in Step 1b above were used for further analysis. These results, which did not include any other information about the rest of the G domain, contained the id of the protein from which the sequence came, the actual sequence and its position in the respective protein. All the results from Step 1b (options i,ii,iii) were merged separately for each G-Protein family. Redundant results, which had the same G5 box sequence and position in a particular protein, were removed (Figure 3c).

For example-

(1) “XXXSAXX” with one mismatch gives following outputs -

| ProteinID | G1-box | Position | G3-box | Position | G4-box | Position | G5-box | Position |
|---|---|---|---|---|---|---|---|---|
| P49411 | GGGAKFK | 83 | DYVK | 131 | NKAD | 180 | IVGSALC | 215 |
| P49411 | GGAKFKK | 84 | DYVK | 131 | NKAD | 180 | IVGSALC | 215 |

(2) “XXXSAXX” with no mismatch gives following outputs –

| ProteinID | G1-box | Position | G3-box | Position | G4-box | Position | G5-box | Position |
|---|---|---|---|---|---|---|---|---|
| P49411 | GGAKFKK | 84 | DYVK | 131 | NKAD | 180 | IVGSALC | 215 |
| Q79GC6 | GGEARGY | 40 | DYVK | 86 | NKAD | 135 | VKGSAKL | 170 |

(3) “XXXAXXX” with no mismatch gives following outputs -

| ProteinID | G1-box | Position | G3-box | Position | G4-box | Position | G5-box | Position |
|---|---|---|---|---|---|---|---|---|
| P02994 | GVTTEVK | 281 | DPPK | 329 | EKND | 374 | CVEAFSE | 408 |
| P05197 | GSGLHGW | 214 | DRYF | 262 | EKLD | 316 | WLPAGDA | 342 |

After merging G5 box results from the three outputs above, we have -

| ProteinID | G5-box | Position |
|---|---|---|
| P49411 | IVGSALC | 215 |
| P49411 | IVGSALC | 215 |
| P49411 | IVGSALC | 215 |
| Q79GC6 | VKGSAKL | 170 |
| P02994 | CVEAFSE | 408 |
| P05197 | WLPAGDA | 342 |

The first three rows are redundant results; so we only consider it once and the final output after Step 2 is:

| ProteinID | G5-box | Position |
|---|---|---|
| P49411 | IVGSALC | 215 |
| Q79GC6 | VKGSAKL | 170 |
| P02994 | CVEAFSE | 408 |
| P05197 | WLPAGDA | 342 |

<u>Step 3 - Prediction of new G5 box consensus sequence</u>

To find the underlying pattern in the merged G5 box sequences, the amino acids in the modified G5 box 7-mer sequences were replaced with 1, 2, 3 and 4 Xs at all the possible positions (Figure 3c). Since, the modified G5 box consensus sequence was only 7 amino acids long, a maximum of 4 amino acids were replaced with X, so as not to lose the underlying pattern completely.

For example, after introducing 1, 2, 3 & 4 Xs in a 7-mer G5 sequence say “IETSAKT”, the following results were obtained:

With 1 X - XETSAKT, IXTSAKT, IEXSAKT, IETXAKT, IETSXKT, IETSAXT, IETSAKX

With 2 Xs - XXTSAKT, XEXSAKT, XETXAKT, XETSXKT, XETSAXT, XETSAKX, IXXSAKT, IXTXAKT, IXTSXKT, IXTSAXT, IXTSAKX, IEXXAKT, IEXSXKT, IEXSAXT, IEXSAKX, IETXXKT, IETXAXT, IETXAKX, IETSXXT, IETSXKX, IETSAXX

With 3 Xs - XXXSAKT, XXTXAKT, XXTSXKT, XXTSAXT, XXTSAKX, XEXXAKT, XEXSXKT, XEXSAXT, XEXSAKX, XETXXKT, XETXAXT, XETXAKX, XETSXXT, XETSXKX, XETSAXX, IXXXAKT, IXXSXKT, IXXSAXT, IXXSAKX, IXTXXKT, IXTXAXT, IXTXAKX, IXTSXXT, IXTSXKX, IXTSAXX, IEXXXKT, IEXXAXT, IEXXAKX, IEXSXXT, IEXSXKX, IEXSAXX, IETXXXT, IETXXKX, IETXAXX, IETSXXX

With 4 Xs- XXXXAKT, XXXSXKT, XXXSAXT, XXXSAKX, XXTXXKT, XXTXAXT, XXTXAKX, XXTSXXT, XXTSXKX, XXTSAXX, XEXXXKT, XEXXAXT, XEXXAKX, XEXSXXT, XEXSXKX, XEXSAXX, XETXXXT, XETXXKX, XETXAXX, XETSXXX, IXXXXKT, IXXXAXT, IXXXAKX, IXXSXXT, IXXSXKX, IXXSAXX, IXTXXXT, IXTXXKX, IXTXAXX, IXTSXXX, IEXXXXT, IEXXXKX, IEXXAXX, IEXSXXX, IETXXXX

After performing the above exercise of replacing amino acids in the modified 7-mer G5 box with Xs, the new G5 box motifs obtained for several proteins per G protein family became identical. These identical motifs were then clustered. For example, let us consider three different 7-mers namely ABCSADE (sequence 1) from protein 1, AFGSAHE (sequence 2) from protein 2, ABDSAGE (sequence 3) from protein 3. After introducing 3 Xs, sequences 1, 2 and 3 become AXXSAXE. These can be clustered together as AXXSAXE: {protein1, protein2, protein3} and we can say that 'AXXSAXE' is a consensus motif for the 3 proteins from the given protein family.

The "coverage" for each predicted consensus G5 motif (with Xs) was calculated as the percentage of proteins from that particular G protein family that had the given motif. Position weighted matrices (PWM) were also calculated for the G5 motifs using ggseqlogo package available on GitHub (43). The sequence logos of predicted G boxes were visualized using the ggseqlogo package, a visualization tool which uses polygons to draw elongated letters. The height of each letter was computed based on their relative frequencies at the specified position.

The G5 motifs which had the highest coverage and were prominent in the PWM were selected as the new consensus G5 box sequence for the given G protein family. Xs at the two ends of 7-mer consensus motifs were trimmed to give more concise G5 box consensus sequence (Suppl. Table 6).

<u>Step 4- G Domain prediction with new G5 box sequences</u>

Step 1 of SMA was repeated to search for all possible G-domains (along with putative G boxes) in each G protein sequence, using the family-specific constraints of consensus sequence, spacers and allowed mismatches listed in Suppl. Table 3 along with the additional constraint of using the new G5 box sequences predicted in Step 3.

### Supplementary Figures

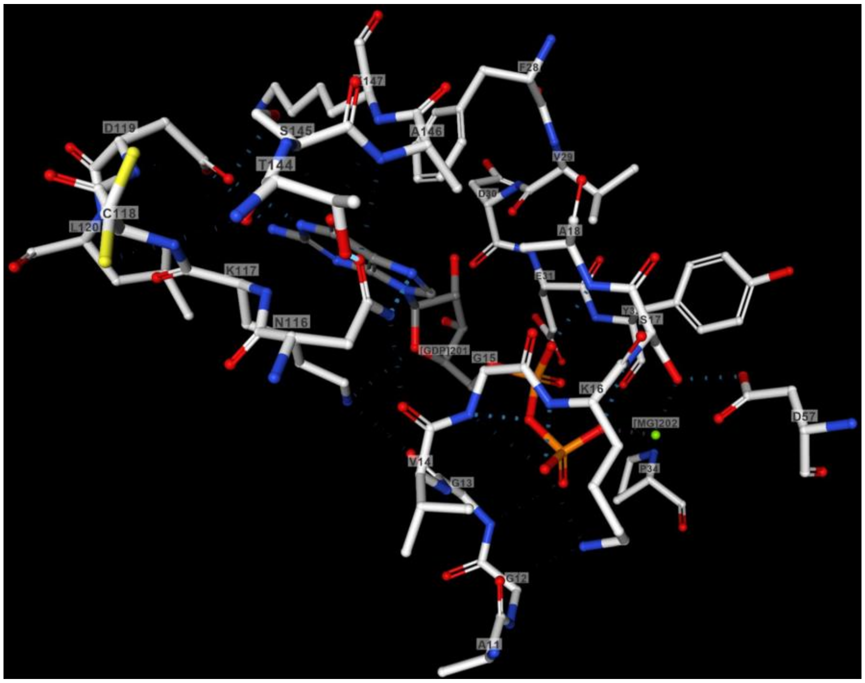


**Suppl. Figure 1: Interactions of GDP with a G protein.** Crystal structure of human NRAS GTPase bound with GDP (PDB id: 3CON) is used to locate the direct contacts of GDP with NRAS. Ligand view option in 3D view tab on the RSCB page used. Blue dashed lines represent the hydrogen bonds between G boxes of NRAS (G1 box: GAGGVG$_{15}$K$_{16}$, G4 box: N$_{116}$KCD$_{119}$ and G5 box: SA$_{146}$K) and GDP.

**Suppl. Figure 2: Comparison of predicted G domain boundaries (in all G protein families), before and after SMA-Step 3**

**Suppl. Figure 3: Comparison of unique G boxes (in all G protein families) predicted before and after SMA-Step 3**

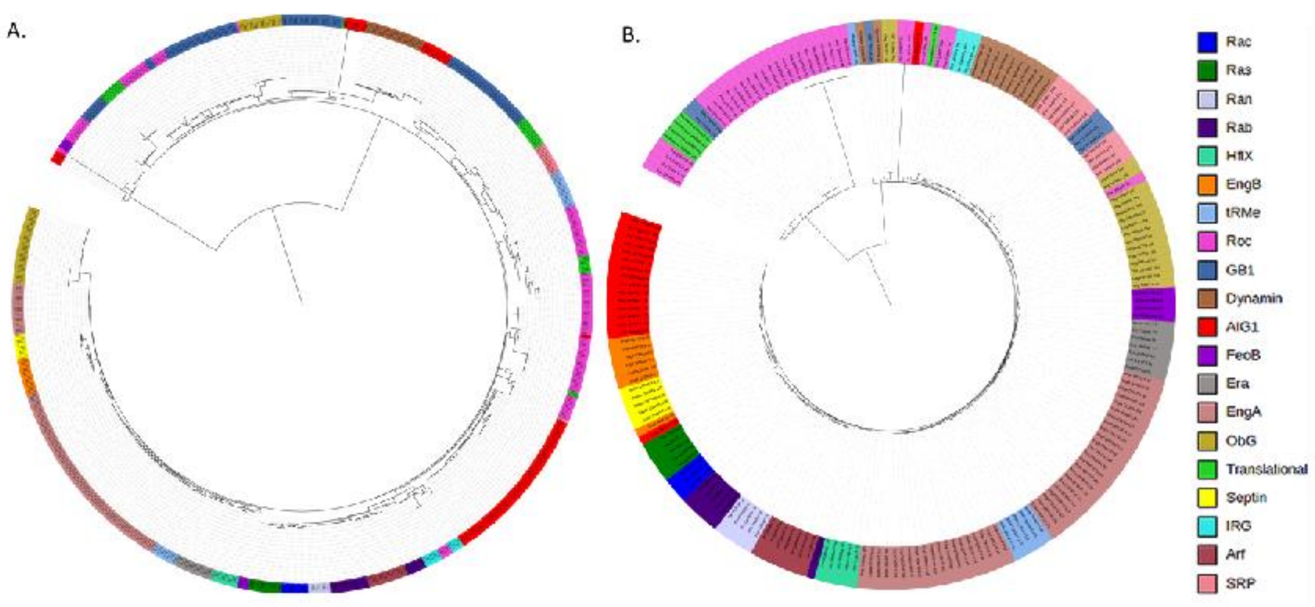


**Suppl. Figure 4: Comparison of phylogenetic trees generated using G domains predicted before and after SMA-Step 3.** Predicted G domains of representative proteins (a) before SMA-Step 3 (b) after SMA-Step 3. Tree scale 1.

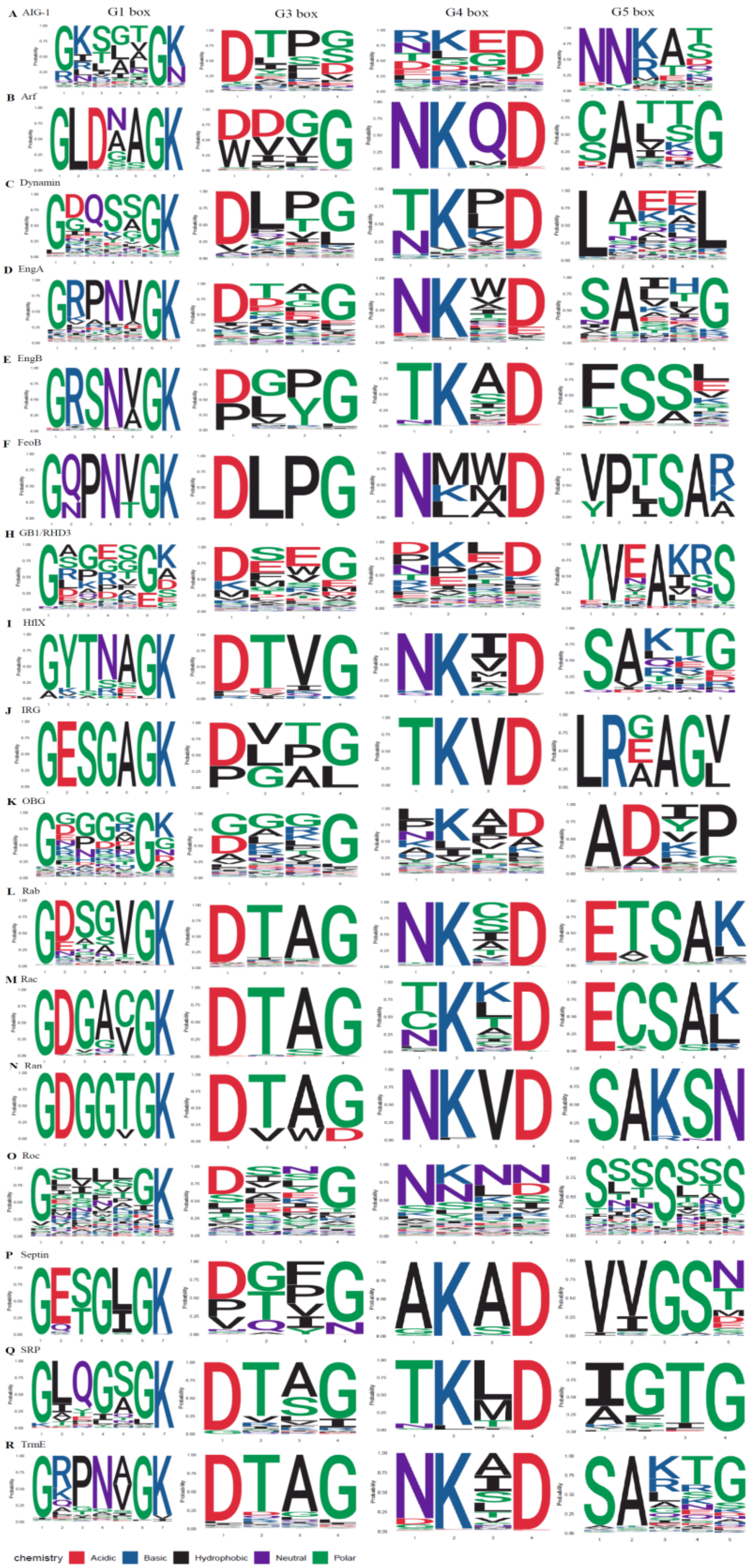


**Suppl. Figure 5: G protein family-specific G box motifs predicted using SMA3**

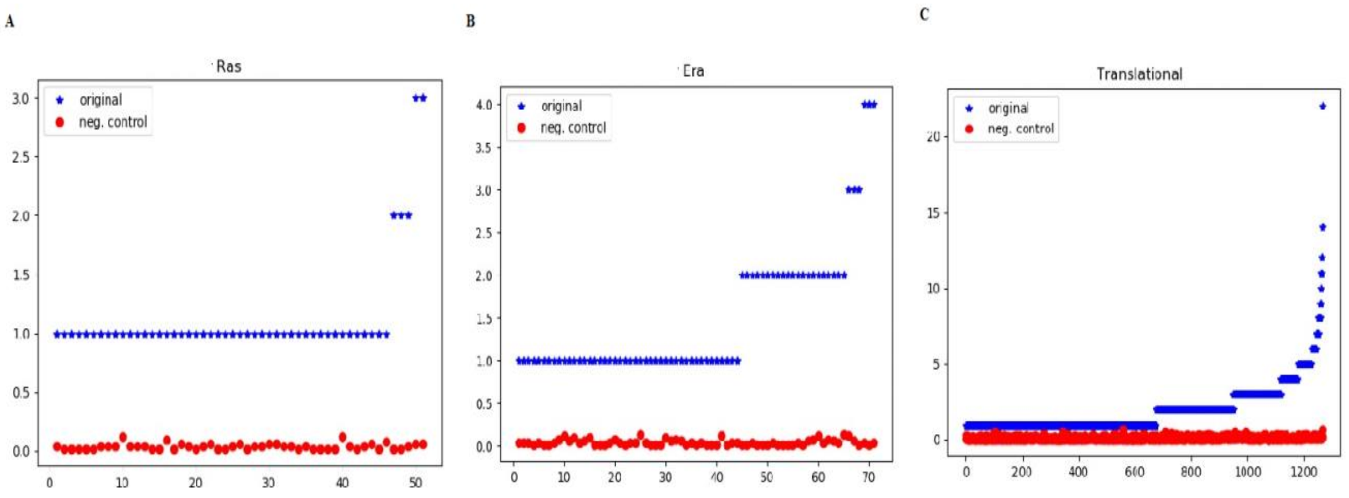


**Suppl. Figure 6. Significance testing of predicted G-domains.** (A) Ras, (B) Era (C) translational families. X-axis=proteins sorted in the order of number of matches obtained; Y-axis= number of predicted G-domains [in actual protein sequence (blue) and average of no. of G-domains identified after 50 shuffles (red)].

**Suppl. Table 1. Difference in the number of input sequences after removal of similarity bias**

| G protein Family | Total no. of protein sequences | No. of protein sequences after removal of similarity bias |
|---|---|---|
| AIG1 | 44 | 44 |
| Arf | 184 | 140 |
| Dynamin | 126 | 115 |
| EngA | 622 | 424 |
| EngB | 548 | 401 |
| Era | 360 | 188 |
| FeoB | 28 | 19 |
| GB1/RHD3 | 122 | 116 |
| HflX | 26 | 26 |
| IRG | 12 | 10 |
| OBG | 864 | 659 |
| Rab | 500 | 470 |

| | | |
|---|---|---|
| Rho/Rac | 163 | 128 |
| Ran | 50 | 29 |
| Ras | 216 | 190 |
| Roc | 20 | 19 |
| Septin | 95 | 86 |
| SRP | 117 | 99 |
| Translational | 4155 | 2870 |
| TrmE | 596 | 454 |

**Suppl. Table 2: Proteins IDs and corresponding PDB IDs for representative members (from each G protein family) for which structural data is available**

| G protein family | PDB IDs | Protein ID |
|---|---|---|
| AIG1 | 1H65 | Q41009 |
| Arf | 1RRF, 1HUR, 1E0S, 1UPT, 1FZQ, 1KSG, 1MOZ | P84079, P84077, P62330, P40616, Q9WUL7, Q9D0J4, P38116 |
| Dynamin | 3W6N,1DYN | O00429, Q05193 |
| EngA | 1MKY | Q9X1F8 |
| EngB | 1SUL, 3PQC | P38424, Q9X1H7 |
| Era | 1EGA | P06616 |
| FeoB | 2WJG | Q57986 |
| GB1/RHD3 | 1DG3, 3Q5D | P32455, Q8WXF7 |
| HflX | 2QTF | Q980M3 |
| IRG | 1TPZ | Q9QZ85 |
| OBG | 1LNZ | P20964 |
| Rab | 4RKF, 1N6H, 2Y8E, 2E9S, 1ZBD, 3QBT, 1T91, 1WMS, | P25228, P20339, O18334, Q9NRW1, P63012, P61006, P51149, P51151 |
| Rho/Rac | 1FOE, 1M7B, 1A2B, 1Z2C | P63000, P61587, P61586, P08134 |
| Ran | 1I2M | P62826 |
| Ras | 2N9C, 2DPX, 3Q85, 2CJW, 1X1R, 121P | P01111, P55042, Q8VEL9, P55040, O08989, P01112 |
| Roc | 2ZEJ | Q5S007 |
| Septin | 2QAG, 2QA5 | Q16181, Q15019 |
| SRP | 1FFH;1JPJ;1JPN;1LS1;1NG1;1O87;1OKK;1RJ9;1RY1;2C03;2C04;2CNW;2FFH;2IY3;2J45;2J46;2J7P;2NG1;2XKV;3NG1;3ZN8 | O07347 |
| Translational | 4ZCI, 4ZKD, 2H5E, 3VQT, 4W2E, 4N3G, 1R5B, 4C0S, 1D2E, 3WBI, | P32132, Q08491, P0A7I4, B8DIL5, Q5SKA7, G0S8G9, O74718, Q71V39, P49410, P39730, |

| | | |
|---|---|---|
| | 1S0U, 2AHO, 1G7R, 1KJZ, 2D74, 3J4J, 1D1N, 1D8T | Q58657, Q980A5, O26359, Q9V1G0, Q8U082, P48515, P04766, P0CE47 |
| TrmE | 1RFL | P25522 |

### Suppl. Table 3: Manually curated spacings between consecutive G boxes and number of mismatches allowed in different G boxes

| G Protein family | Putative amino acid spacers between G1 and G3 box | Putative amino acid spacers between G3 and G4 box | Putative amino acid spacers between G4 and G5 box | Number of mismatches allowed in G1 box consensus sequence (GXXXGK) | Number of mismatches allowed in G3 box consensus sequence (DXXG) | Number of mismatches allowed in G4 box consensus sequence (NKXD) | Number of mismatches allowed in G5 box consensus sequence (SAX) |
|---|---|---|---|---|---|---|---|
| AIG-1 | 45-55 | 65-75 | 30-40 | 1 | 1 | 2 | 1 |
| Arf | 40-50 | 55-65 | 25-35 | 1 | 1 | 1 | 1 |
| Dynamin | 95-105<br>95-105 | 65-75<br>55-75 | 25-35<br>40-50/65-80. | 1 | 1 | 1 | 1 |
| EngA | 40-80 | 40-80 | 20-80 | 1 | 1 | 1 | 1 |
| EngB | 40-45<br>40-50 | 65-75<br>65-75 | 20-35<br>20-35 | 1 | 1 | 1 | 1 |
| Era | 40-50 | 60-70 | 25-35 | 1 | 1 | 1 | 1 |
| FeoB | 45-50 | 60-65 | 20-30 | 1 | 1 | 1 | 1 |
| GB1/RHD3 | 55-75 | 65-85 | 40-70 | 1 | 1 | 2 | 1 |
| Hflx | 40-50<br>60-70 | 65-75<br>130-140 | 20-35<br>20-35 | 1 | 1 | 1 | 1 |
| IRG | 45- 55 | 55-65 | 30-40 | 1 | 1 | 1 | 1 |
| OBG | 45-50<br>65-70 | 60-75<br>130-140 | 20-30<br>25-35 | 1 | 1 | 2 | 1 |
| Rab | 45-60 | 55-65 | 25-35 | 1 | 1 | 1 | 1 |
| Ran | 45-50 | 55-65 | 25-35 | 1 | 1 | 1 | 1 |
| Ras | 45-50 | 55-65 | 25-35 | 1 | 1 | 1 | 1 |
| Rho/Rac | 45-50 | 55-65 | 40-50 | 1 | 1 | 1 | 1 |
| Roc | 45-55 | 55-65 | 35-45 | 1 | 1 | 2 | 1 |
| Septin | 50-60 | 75-85 | 50-60 | 1 | 1 | 1 | 1 |
| SRP | 80-90 | 55-65 | 20-30 | 1 | 1 | 1 | 1 |
| Translational | 45-50<br>45-50 | 45-55<br>60-80 | 25-35<br>30-45 | 1 | 1 | 1 | 1 |

| | | | | | | | |
|---|---|---|---|---|---|---|---|
| | 60-65<br>65-70 | 50-60<br>50-60 | 60-70<br>110-120 | | | | |
| TrmE | 45-50 | 60-75 | 20-30 | 1 | 1 | 1 | 1 |

**Suppl. Table 4: Examples of proteins from the same family which had significantly different inter G box spacers**

| | | | | Inter G box spacer | | |
|---|---|---|---|---|---|---|
| **G protein family** | **Protein name** | **Organism** | **Protein structure PDB ID** | **G1-G3** | **G3-G4** | **G4-G5** |
| Translational | Translation initiation factor IF-2 | Bacillus stearothermophilus | 1D1N;1Z9B;2LKC;2LKD;2NBG | 46 | 54 | 34 |
| Translational | Eukaryotic translation initiation factor 5B (eIF-5B) | Saccharomyces cerevisiae | 3WBI;3WBJ;3WBK;4N3S;4NCF;4V8Y;4V8Z | 64 | 54 | 66 |
| Translational | Peptide chain release factor 3 (RF-3) | Desulfovibrio vulgaris | 3VQT;3VR1 | 68 | 54 | 115 |
| Dynamin | Mitofusin-1 | Homo sapiens | 5GNR;5GNS;5GNT;5GNU;5GO4;5GOE;5GOF;5GOM;5YEW | 96 | 59 | 47 |
| Dynamin | Dynamin-1 | Homo sapiens | 1DYN;2DYN;2X2E;2X2F;3SNH;3ZYC;3ZYS;4UUD;4UUK;5D3Q;6DLU;6DLV | 98 | 69 | 28 |

**Suppl. Table 5: Comparison of the number of proteins generating SMA output using "SAX" or the SMA3-predicted motif as G5 box sequence**

| **G Protein Family** | **Output upon using "SAX" as G5 box motif** | | **Output upon using SMA3-predicted G5 box motif** | |
|---|---|---|---|---|
| | **Number of proteins** | **Percentage of proteins** | **Number of proteins** | **Percentage of proteins** |
| AIG1 | 42 | 95.4 | 27 | 61.3 |
| Arf | 130 | 92.8 | 102 | 72.8 |
| Dynamin | 100 | 86.9 | 91 | 79.1 |
| EngA | 424 | 100 | 424 | 100 |
| EngB | 373 | 93 | 304 | 75.8 |
| Era | 174 | 92.5 | 154 | 81.9 |

| | | | | |
|---|---|---|---|---|
| FeoB | 14 | 77.7 | 4 | 22.2 |
| GB1 | 116 | 100 | 58 | 50 |
| Hflx | 26 | 100 | 20 | 76.9 |
| IRG | 4 | 40 | 3 | 30 |
| OBG | 658 | 99.8 | 613 | 93.01 |
| Rab | 442 | 94.04 | 424 | 90.21 |
| Rho/Rac | 101 | 78.9 | 63 | 49.2 |
| Ran | 28 | 96.5 | 28 | 96.5 |
| Ras | 170 | 89.4 | 168 | 88.4 |
| Roc | 19 | 100 | 18 | 94.7 |
| Septin | 77 | 89.5 | 68 | 79.06 |
| SRP | 42 | 42.4 | 44 | 44.4 |
| Translational | 2562 | 89.2 | 1433 | 49.9 |
| TrmE | 372 | 81.9 | 230 | 50.6 |

**Suppl. Table 6: Predicted G5 box sequence for all twenty G protein families**

| G protein family | G5 box motif identified after SMA analysis |
|---|---|
| AIG-1 | NNXAX |
| Arf | CAXXG |
| Dynamin | LAXXL |
| EngA | SAXXG |
| EngB | FSSX |
| Era | PXSAX |
| FeoB | PPXSAX |
| GB1 or RHD3 | YVXAXXS |
| Hflx | SAXXG |
| IRG | LRXAAX |
| OBG | ADXP |
| Rab | EXSAX |
| Rac/Rho | ECSAX |
| Ran | SAXXN |
| Ras | EXSAX |

| | |
|---|---|
| Roc | SXXSXXS |
| Septin | VXGSX |
| SRP | AXTG |
| Translational | VXXSAX |
| TrmE | SAXXG |

**Suppl. Table 7: Percent reduction in total number of unique G boxes (sum of G1, G3, G4 and G5) predicted per family using the SMA3-generated G5 box sequence**

| G protein family | Percent reduction in total number of unique G boxes predicted |
|---|---|
| AIG-1 | 76.65 |
| Arf | 52.86 |
| Dynamin | 48.16 |
| EngA | 20.39 |
| EngB | 55.94 |
| Era | 34.4 |
| FeoB | 67.75 |
| GB1/RHD3 | 89.8 |
| Hflx | 50.61 |
| IRG | 68 |
| OBG | 54.04 |
| Rab | 48.83 |
| Rho/Rac | 57.7 |
| Ran | 0 |
| Ras | 28.23 |
| Roc | 64.97 |
| Septin | 51.05 |
| SRP | 60.32 |
| Translational | 49.1 |
| TrmE | 46.55 |

**Suppl. Table 8: Comparison of percentage of proteins per G protein family that have the indicated spacer value, before and after SMA3**

| Spacers between G boxes | Comparison of values obtained using SMA3-predicted G5 box sequence versus using "SAX" as G5 box sequence | |
|---|---|---|
| | Similar | Different |
| G1-G3 box | AIG1, Arf, Dynamin, EngA, EngB, Era, GB1, IRG, OBG, Rab, Rac,Ran,Ras, Roc, Septin, translational, TrmE | Hflx, FeoB and SRP |
| G3-G4 box | AIG1, Arf, Dynamin, EngA, EngB, Era, GB1, HflX, IRG, OBG, Rab, Rac, Ran, Ras, Roc, Septin, translational, TrmE | FeoB, SRP |
| G4-G5 box | Dynamin, EngA, FeoB, GB1, Roc, translational | AIG1, Arf, EngB, Era, IRG, HflX, OBG, Rab, Rac, Ran, Ras, Septin, SRP, TrmE |

S**uppl. Table 9: Protein IDs of of alpha subunits of heterotrimeric G protein families used in the study to validate the SMA output**

| **Alpha subunit family** | **Protein ID or PDB ID** | **Output from the code includes the structurally verified sequence** |
|---|---|---|
| Gq | NP_032165.3 | Yes |
| Gi | NP_037277.1 | Yes |
| Gs | 1AZT | Yes |
| G12/13 | NP_034432.1 | Yes |
