## Supplementary figures and images for "G-domain prediction across the diversity of G protein families"

### AIG1.pdf

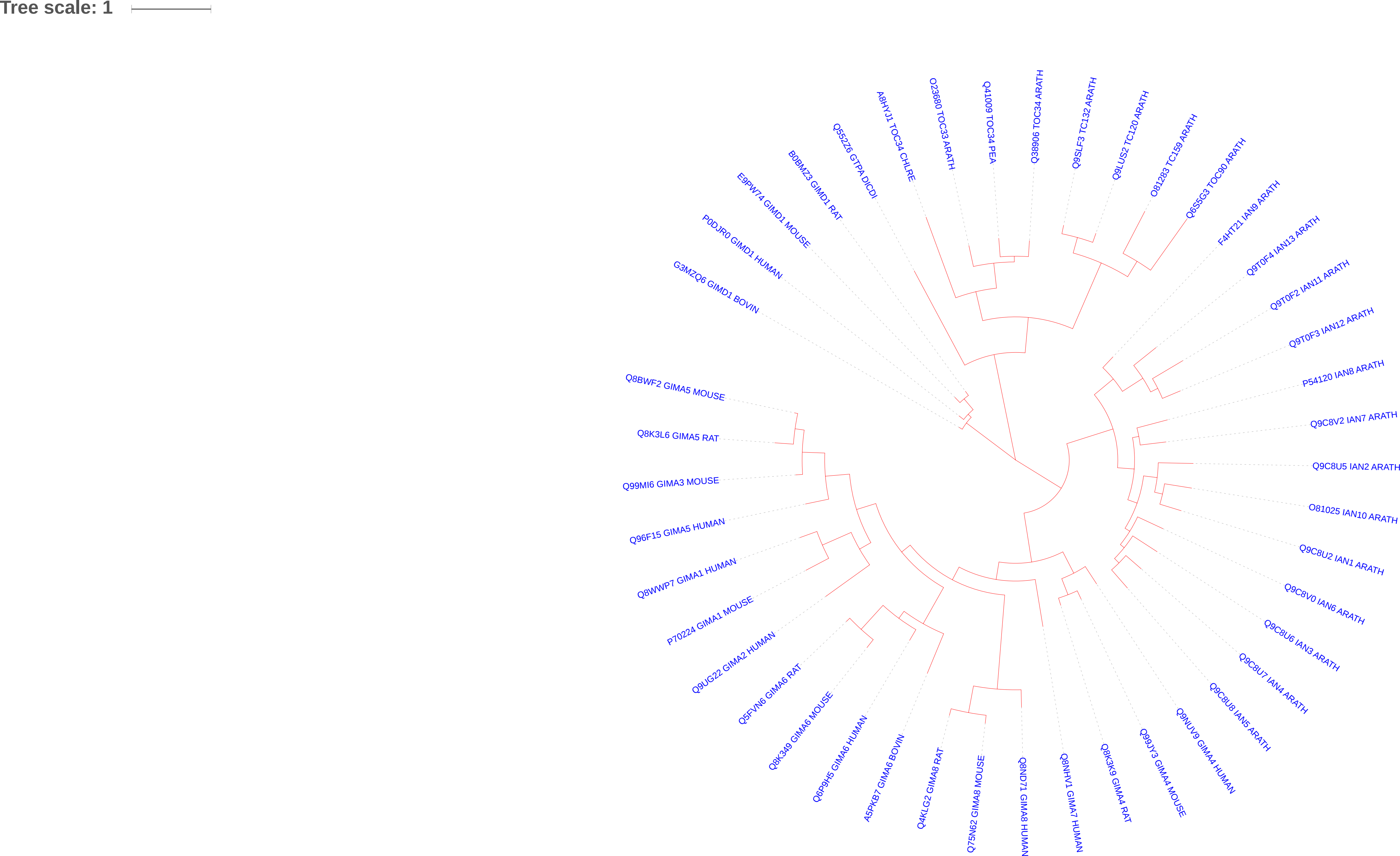

### Arf.pdf

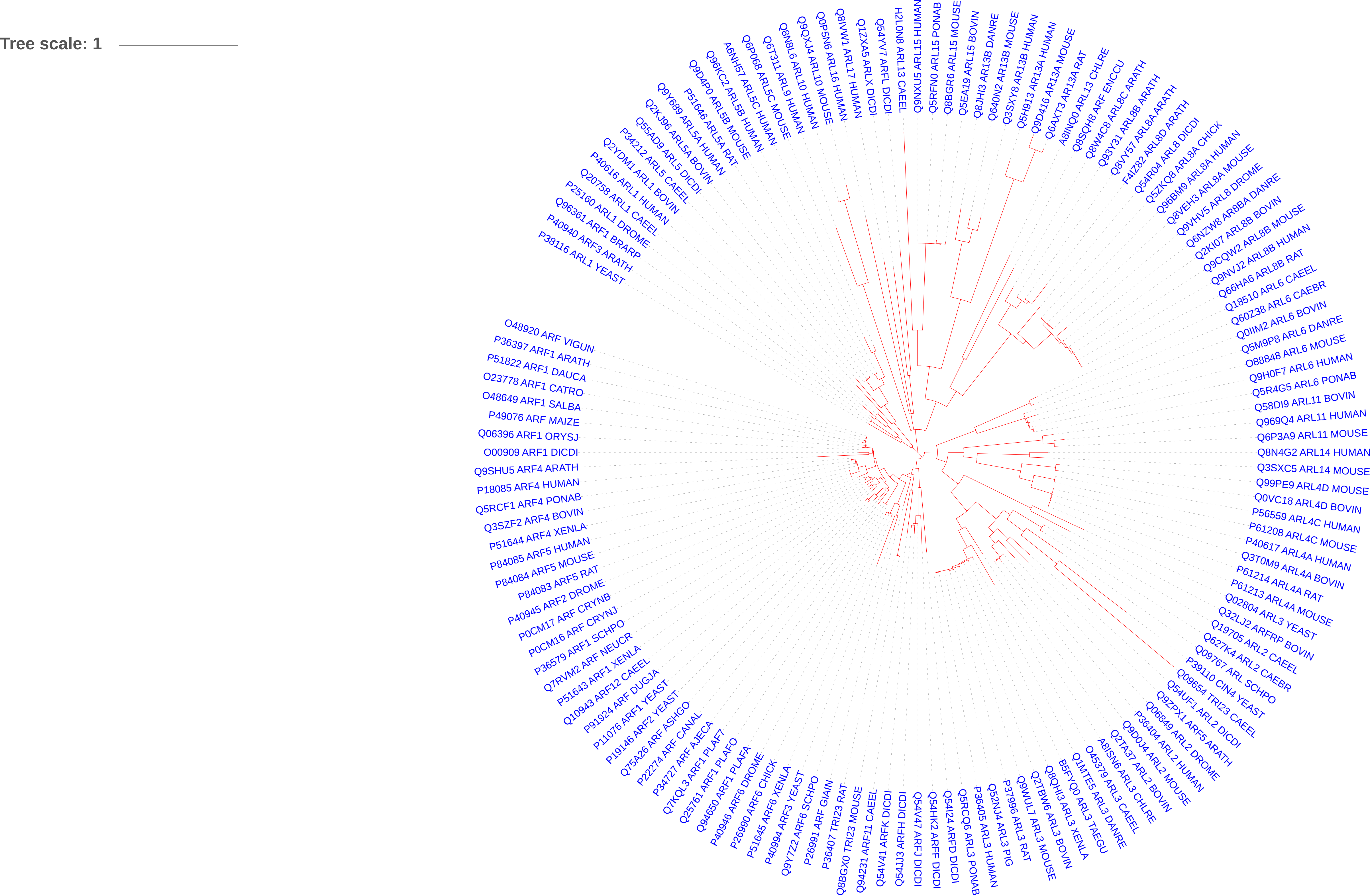

### Dynamin.pdf

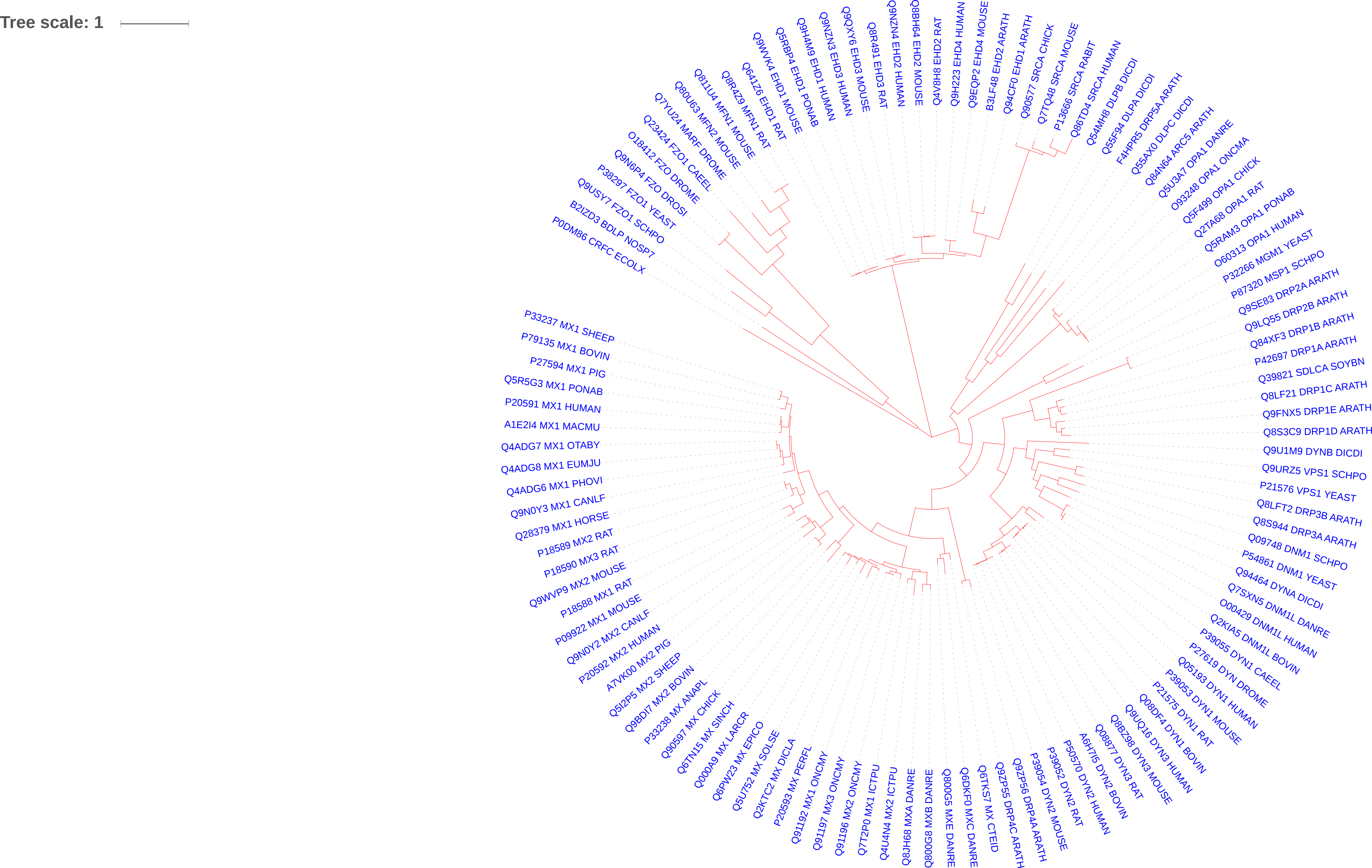

### EngA.pdf

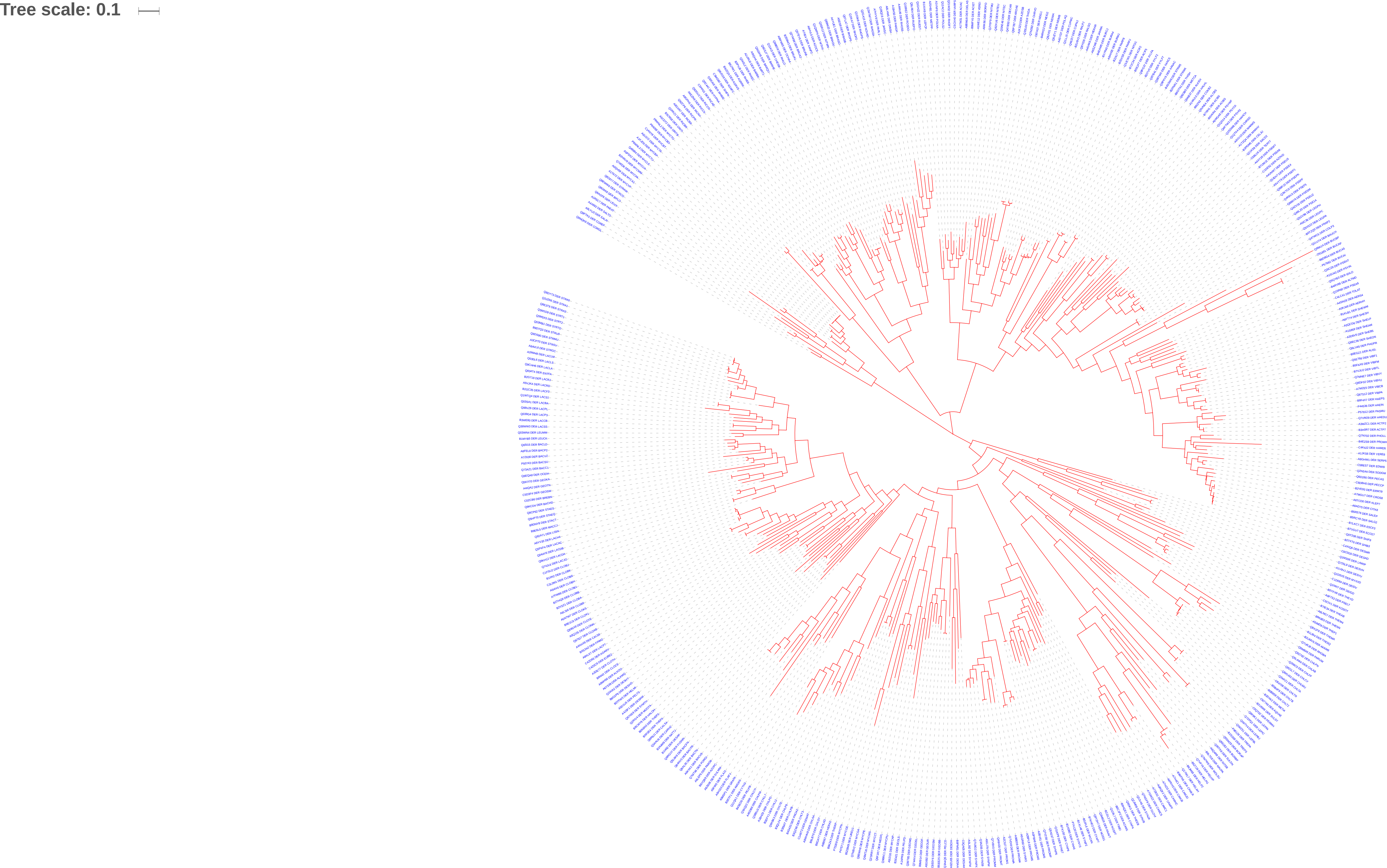

### EngB.pdf

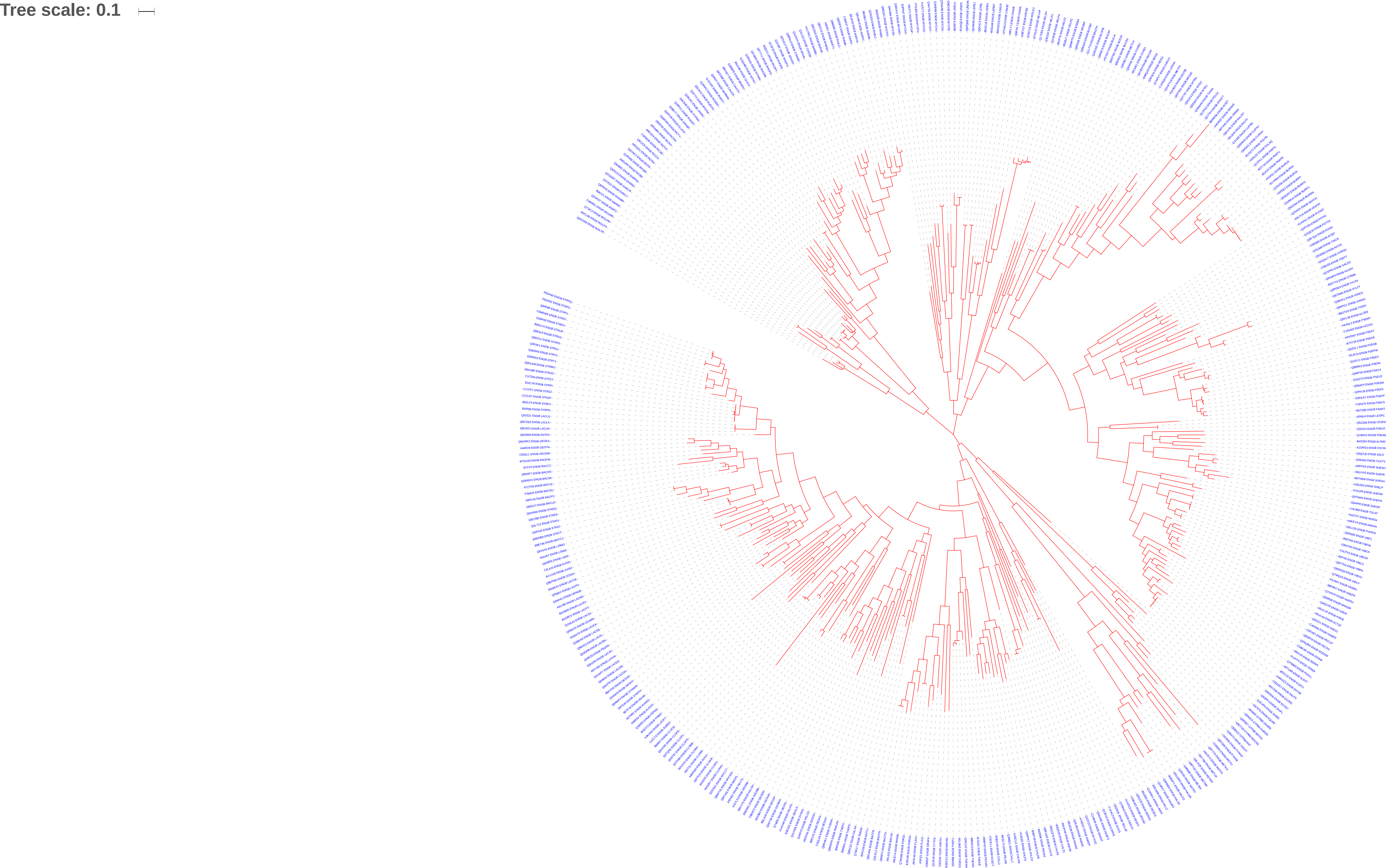

### Era.pdf

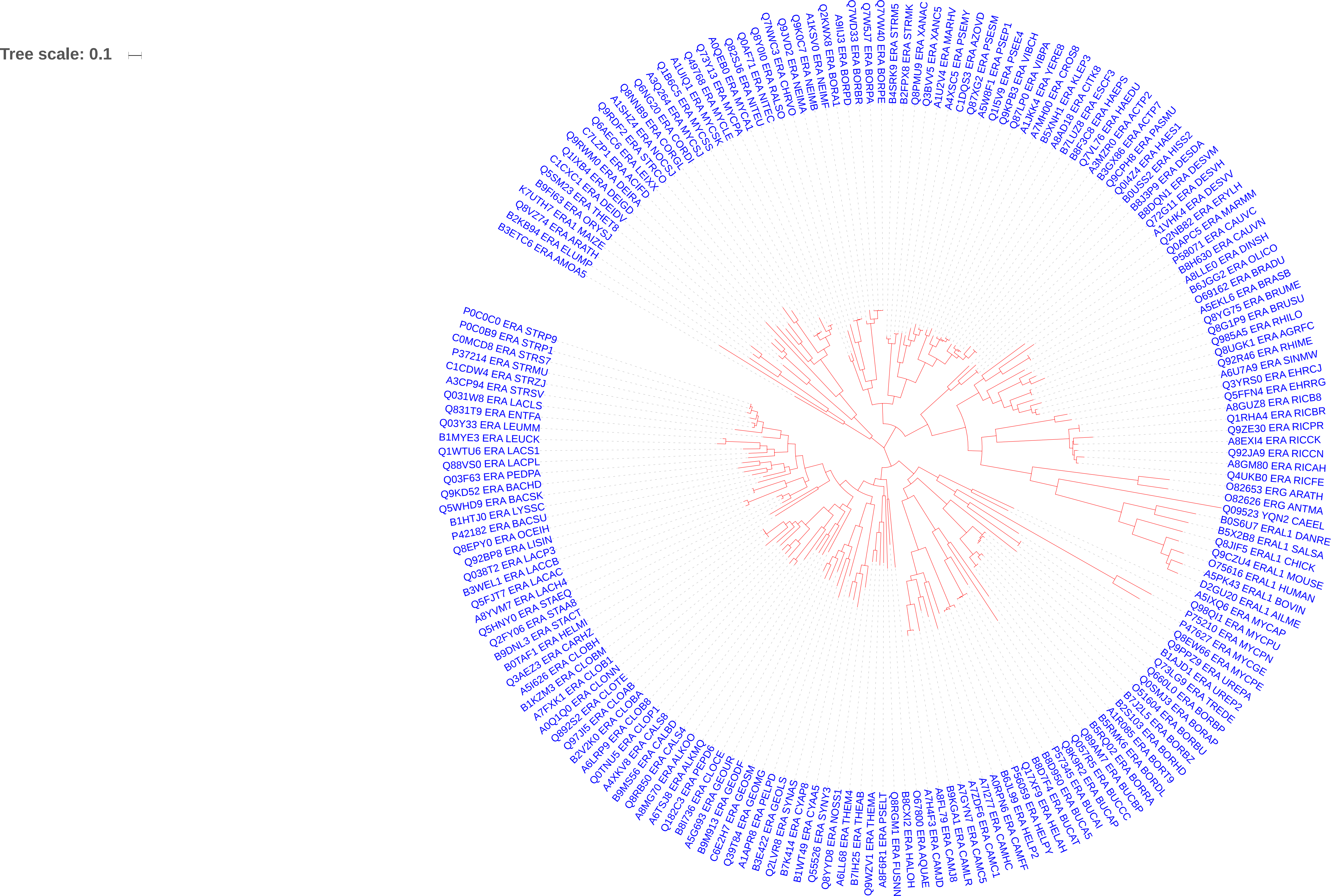

### FeoB.pdf

Tree scale: 0.1

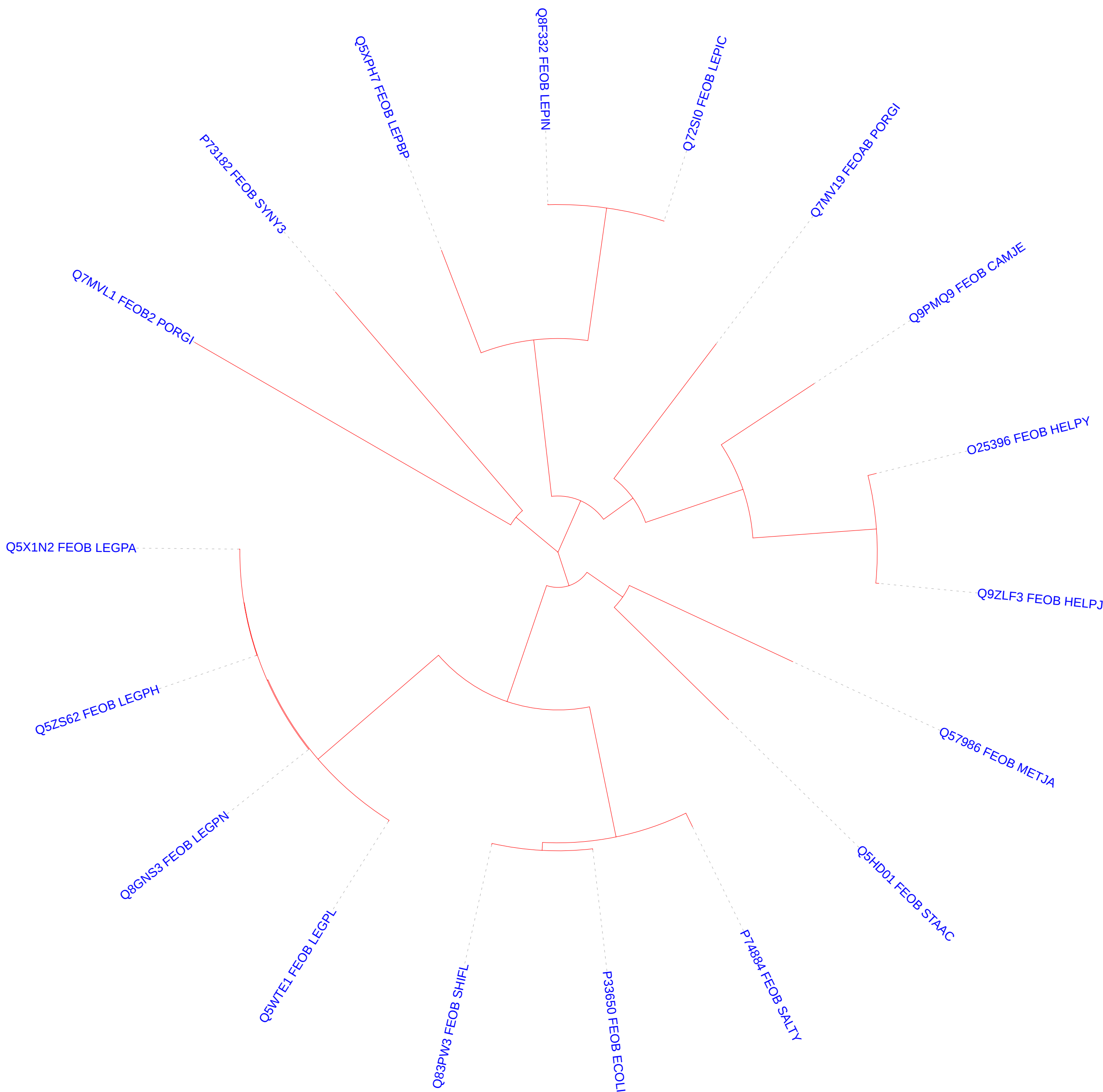

### GB1.pdf

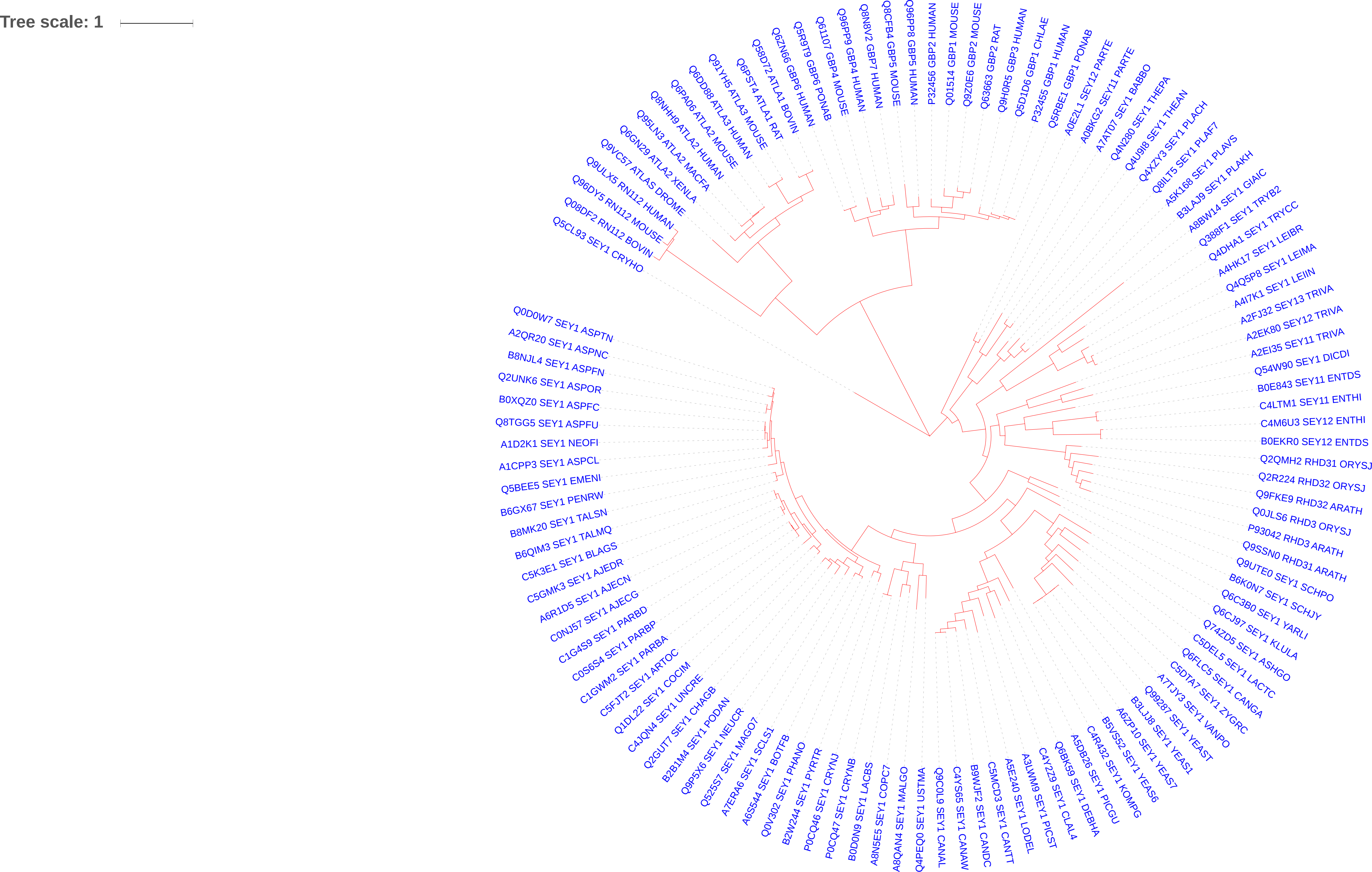

### HflX.pdf

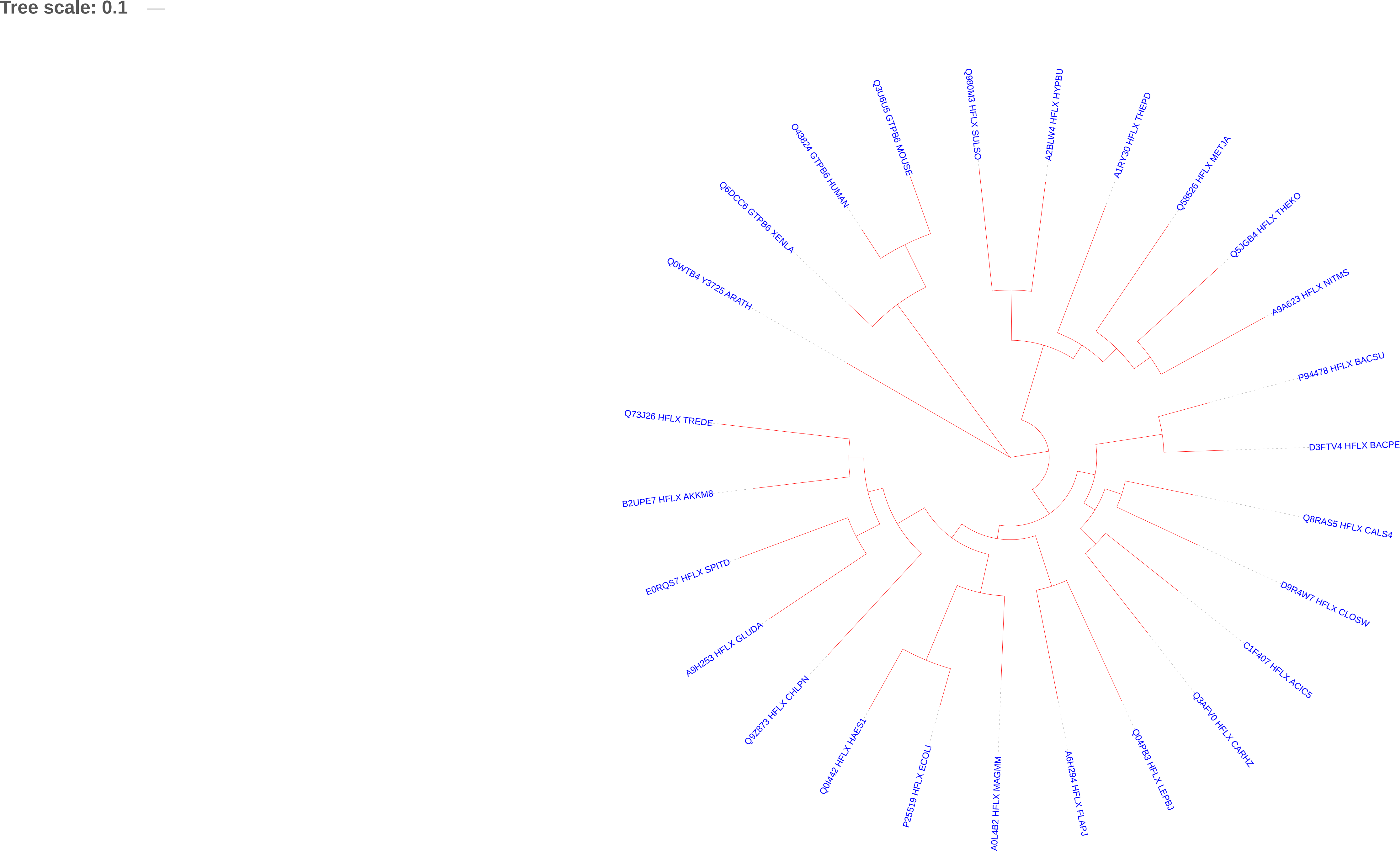

### IRG.pdf

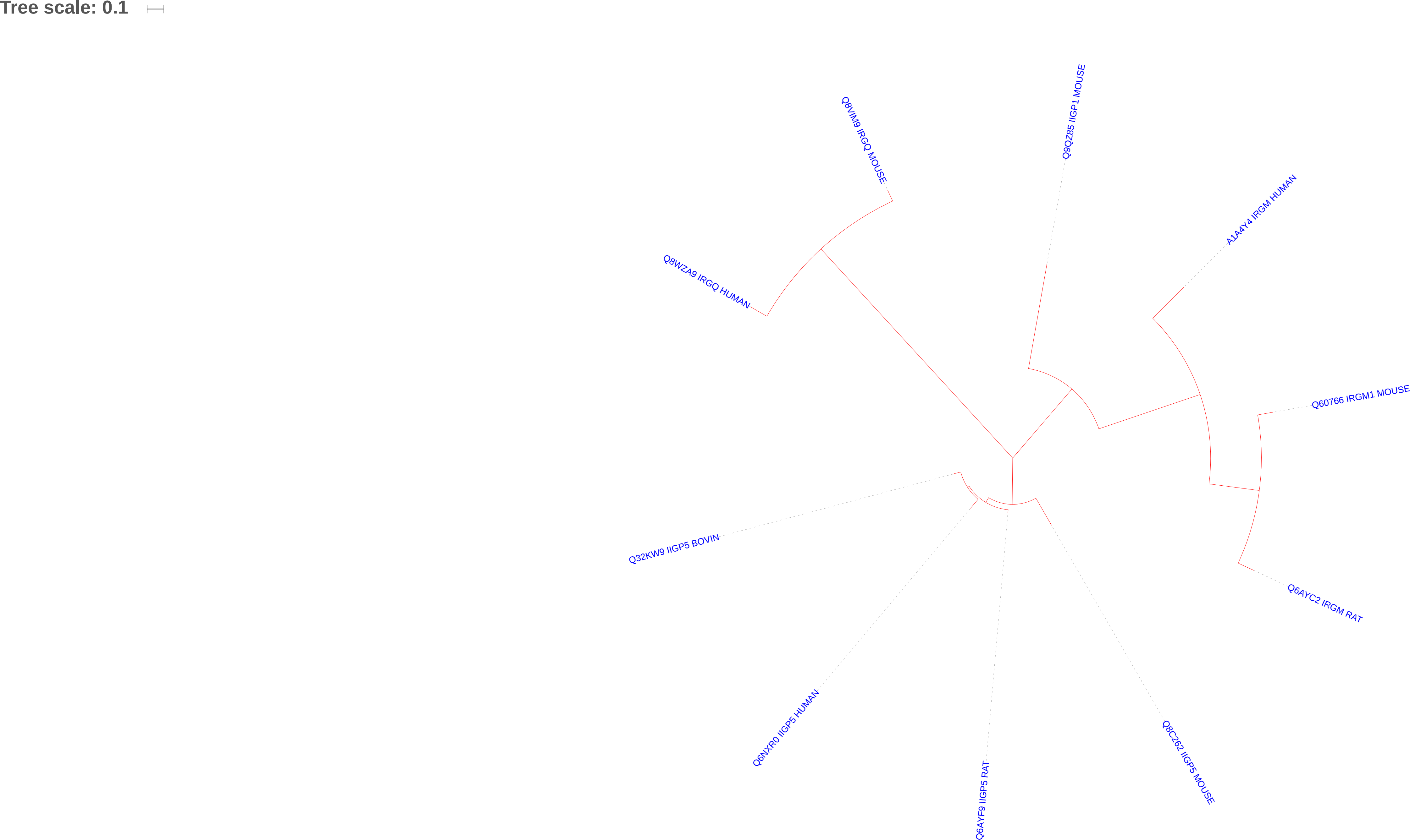

### ObG.pdf

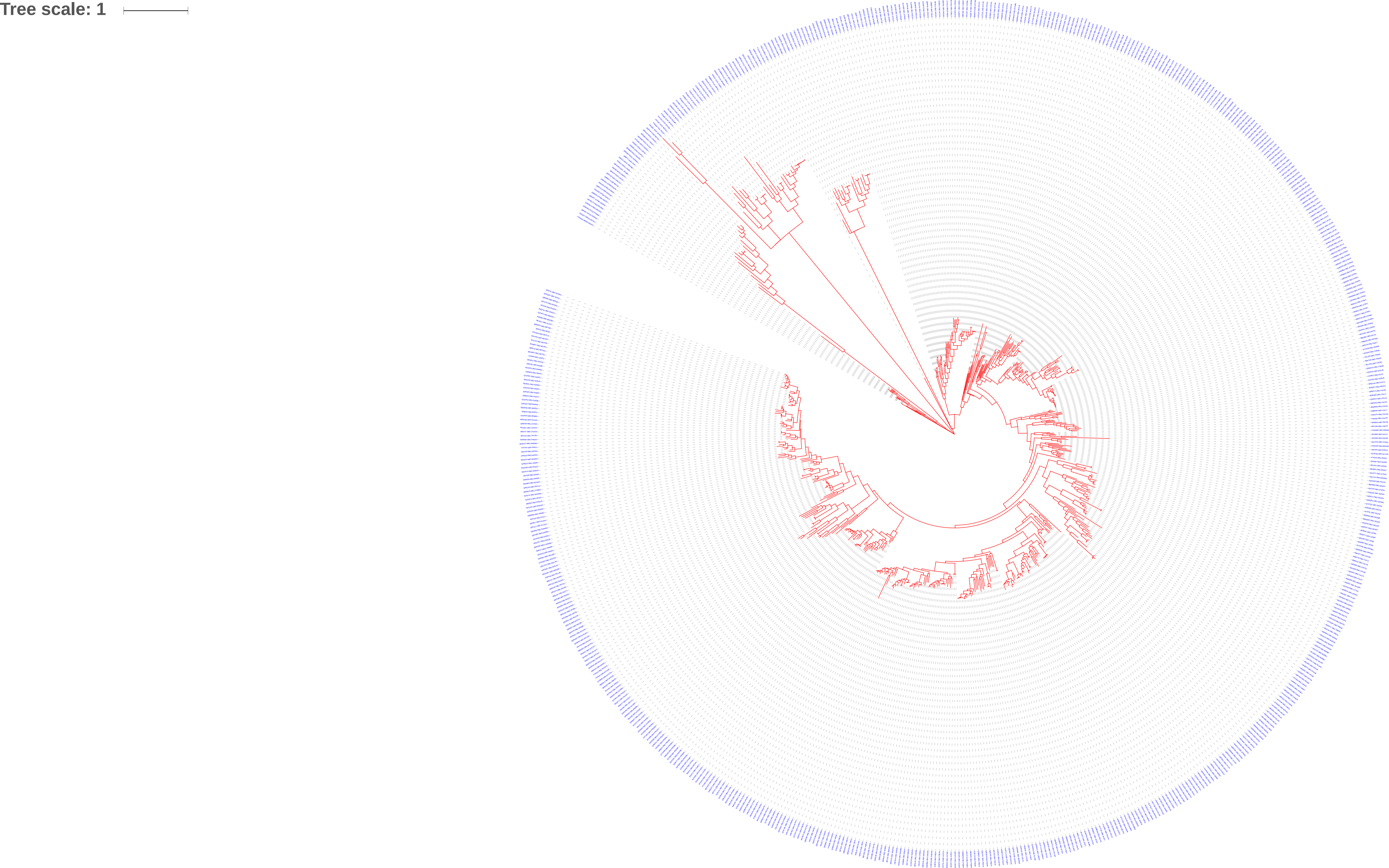

### Rab.pdf

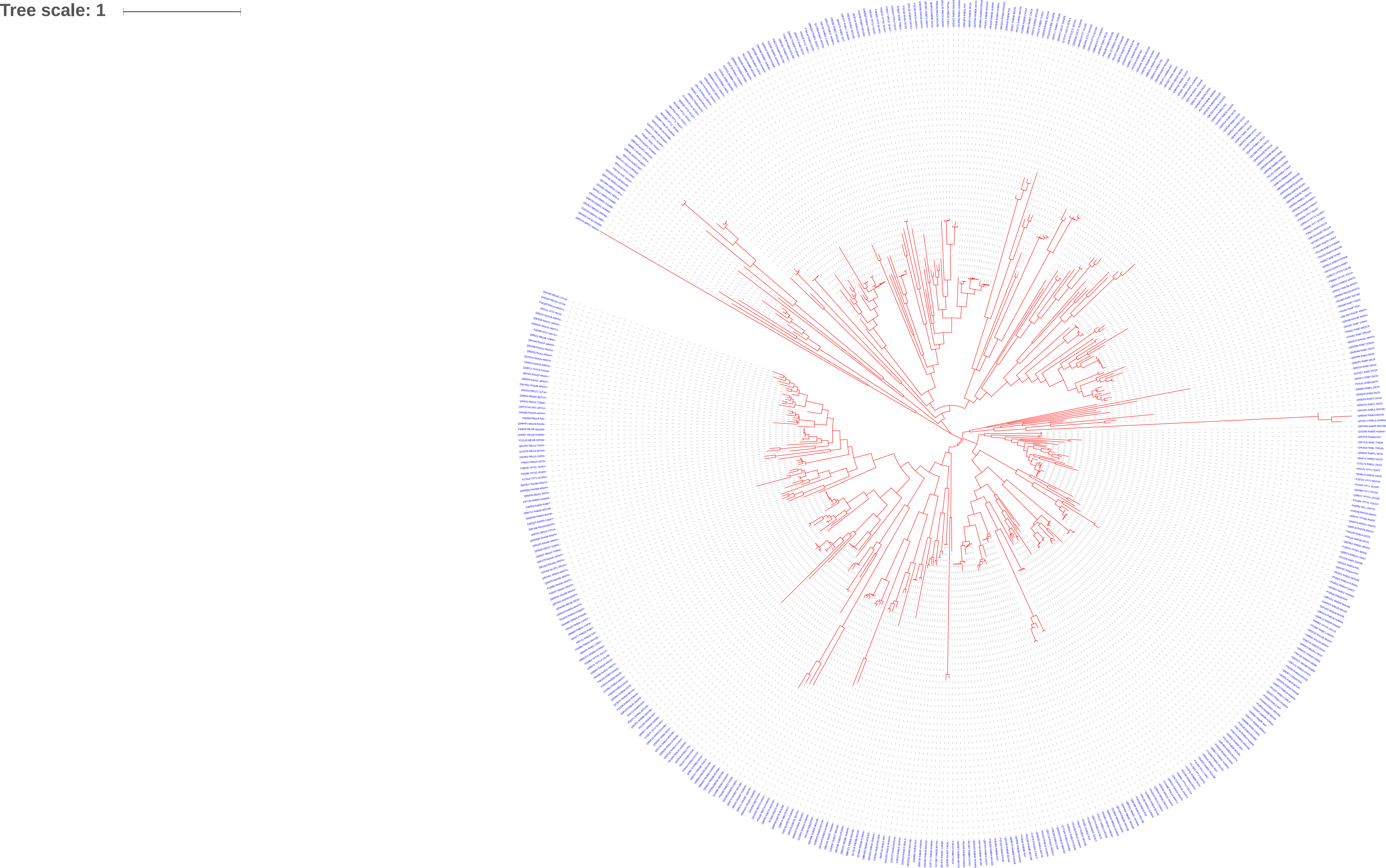

### Rac.pdf

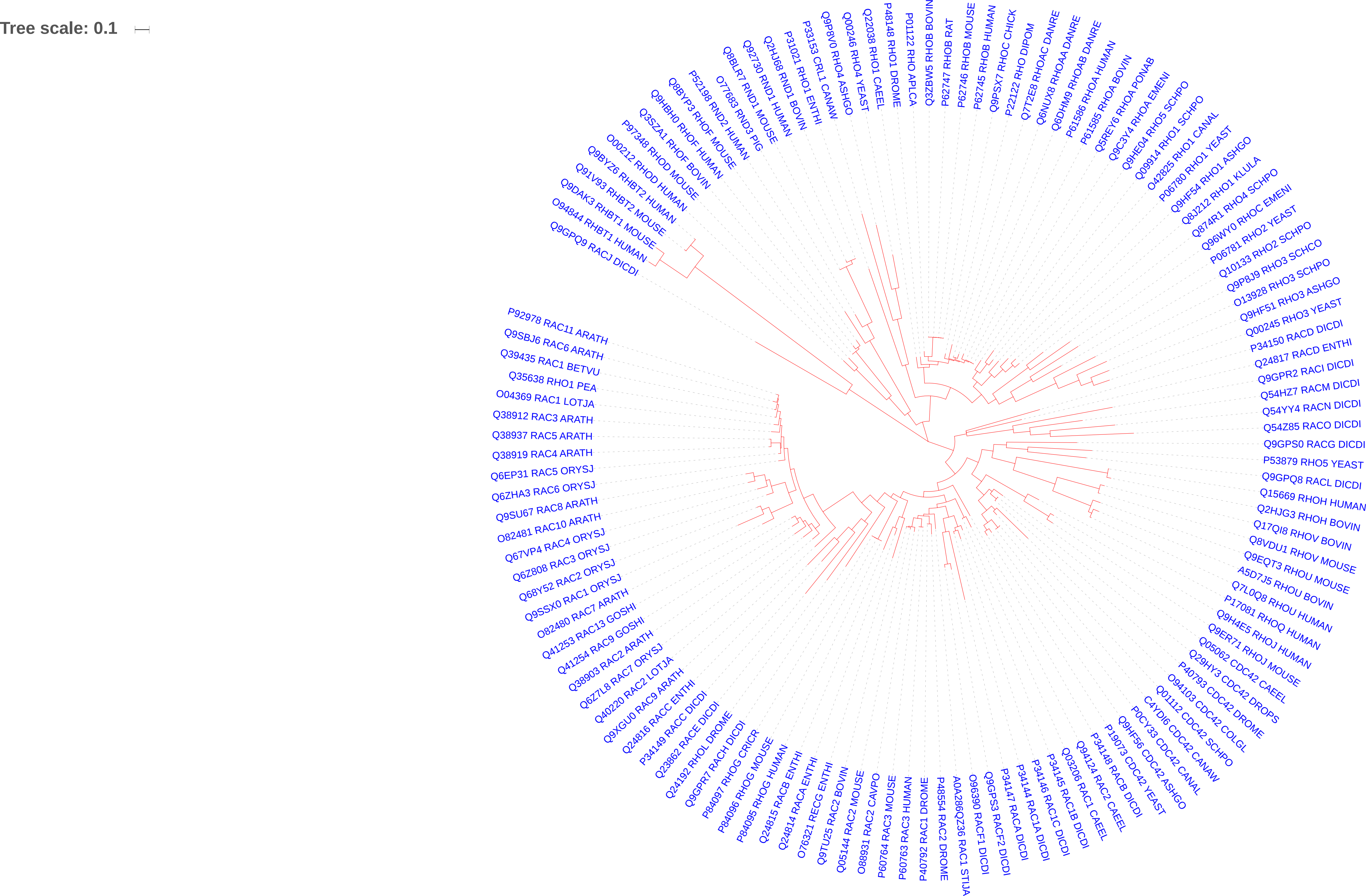

### Ran.pdf

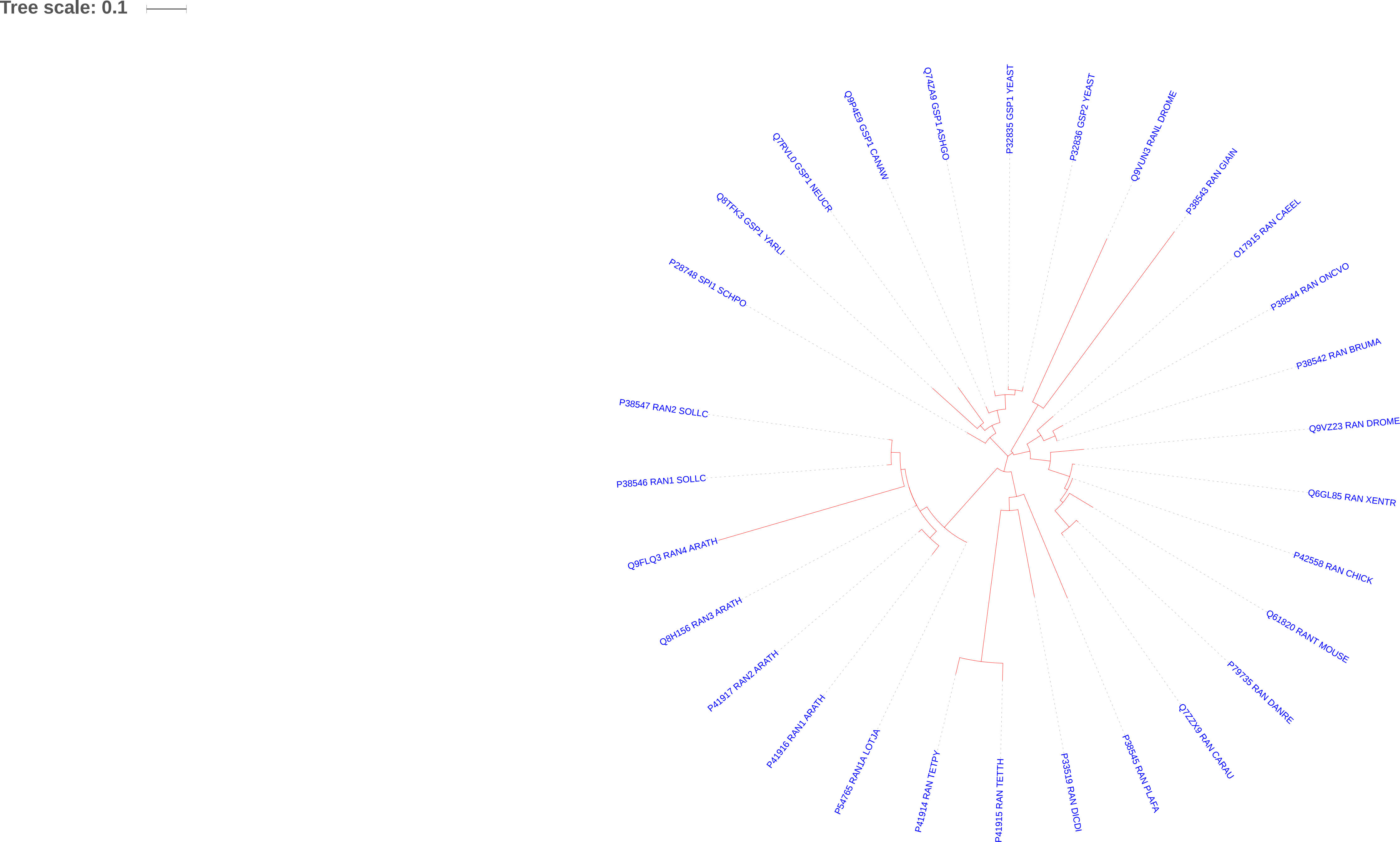

### Ras.pdf

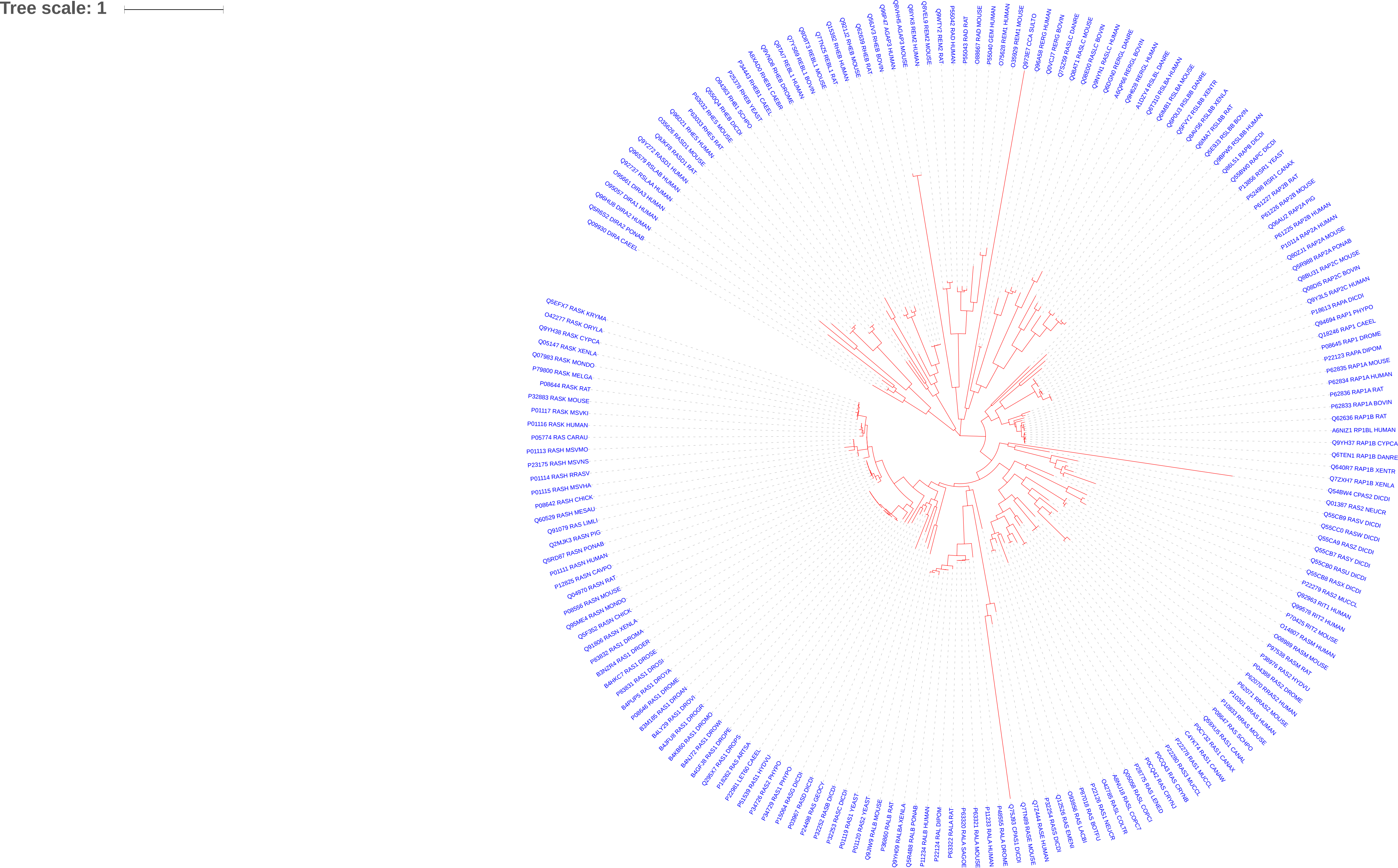

### Roc.pdf

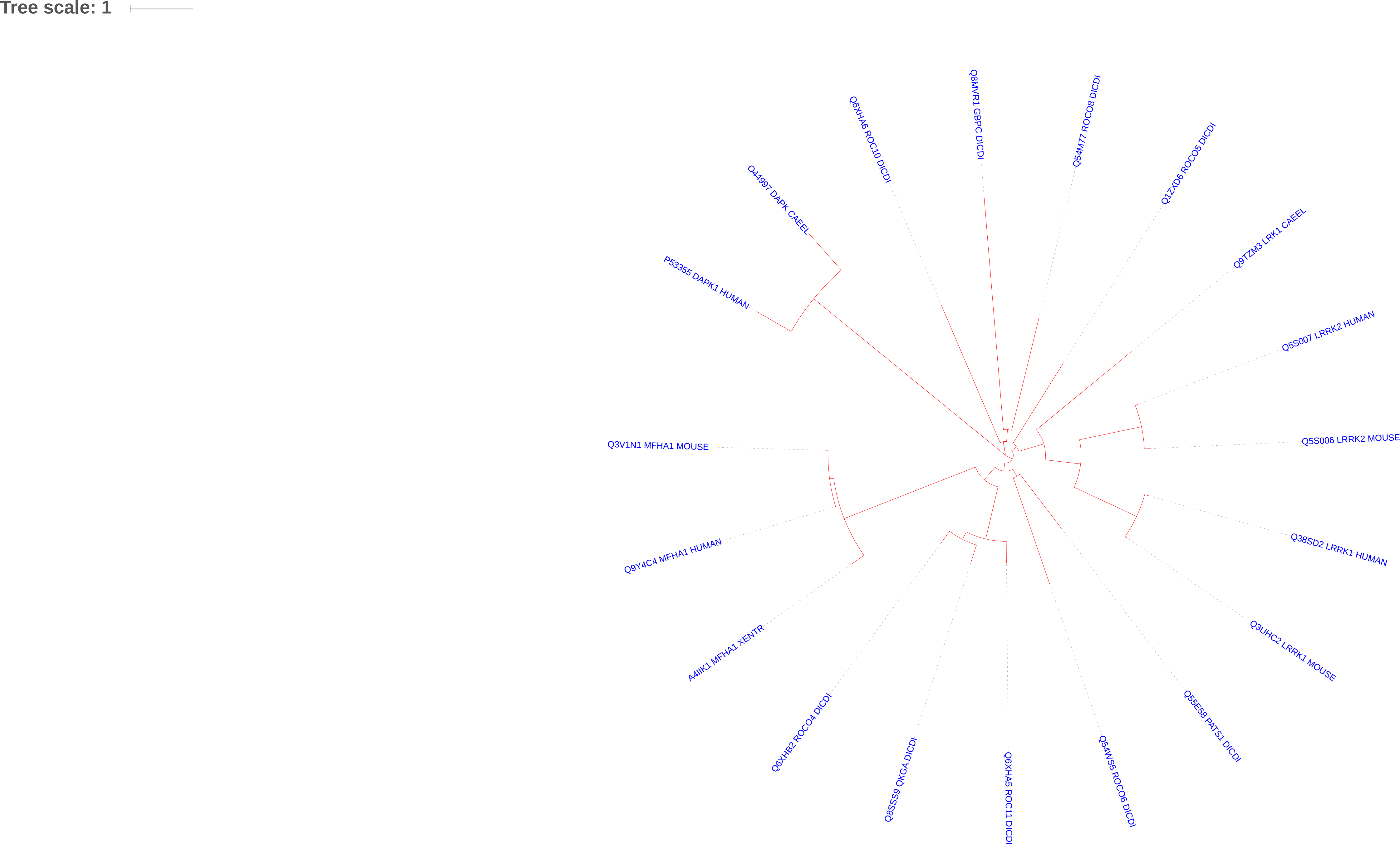

### Septin.pdf

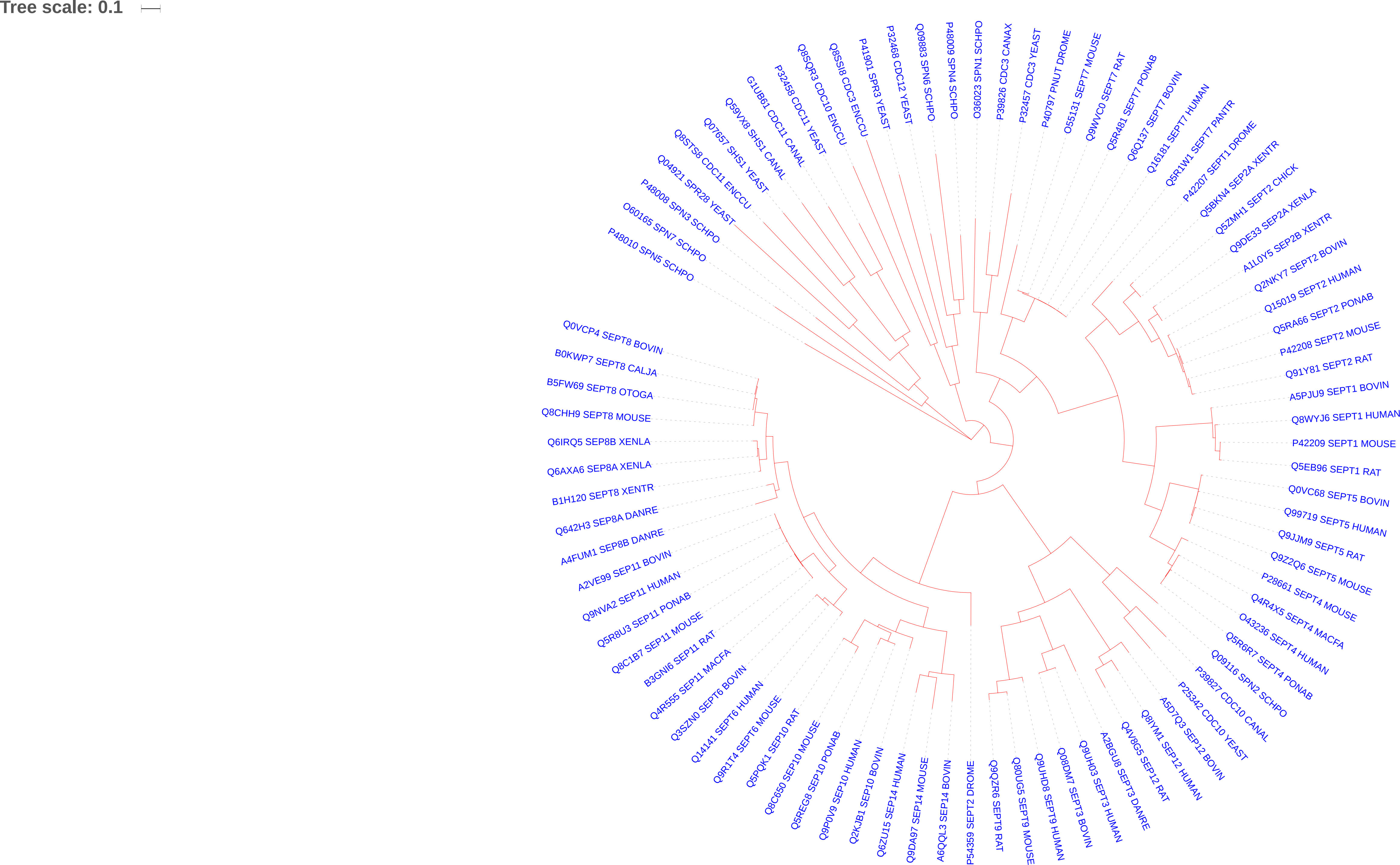

### SRP.pdf

Tree scale: 0.1

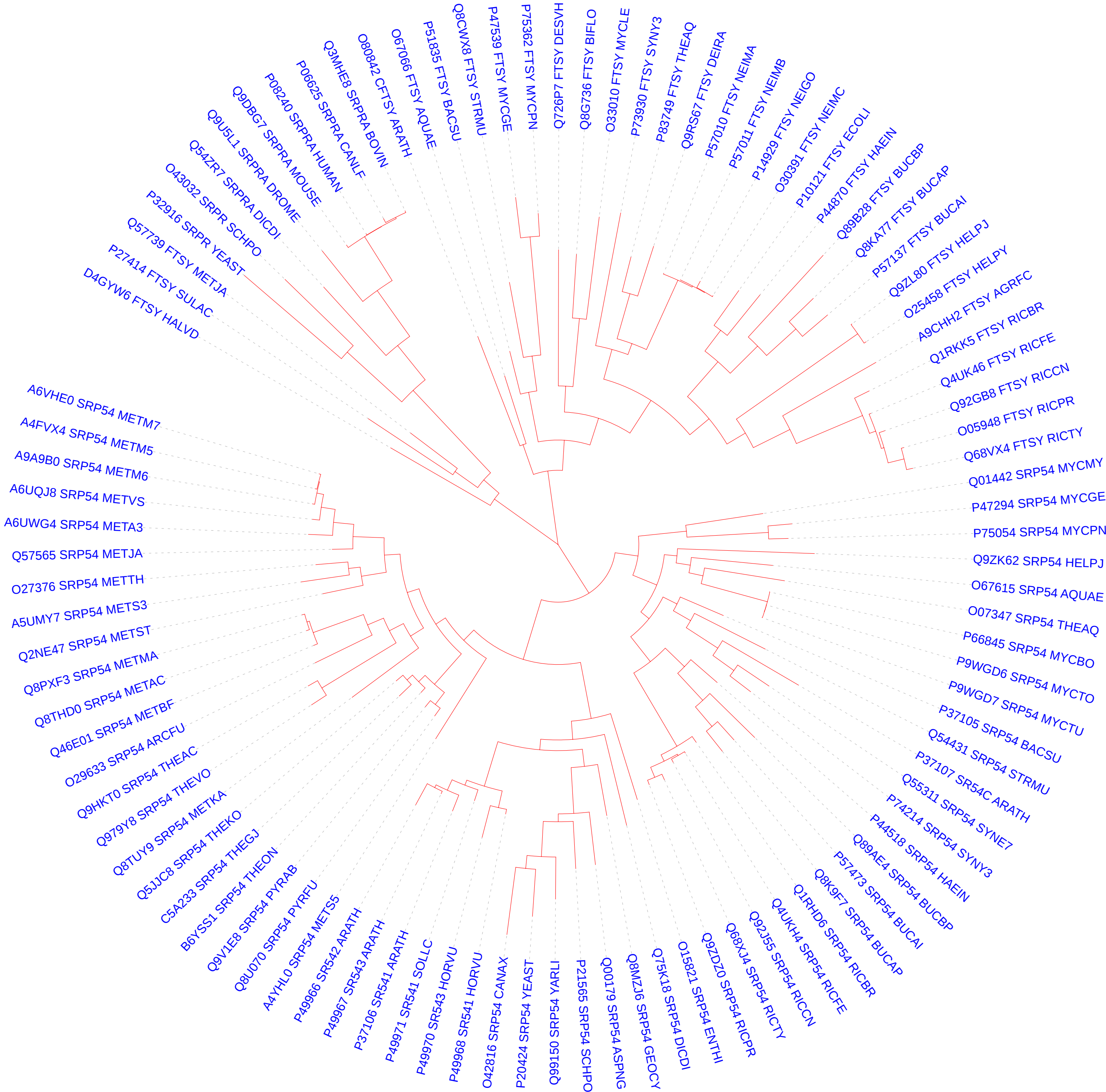

### translational.pdf

Tree scale: 1

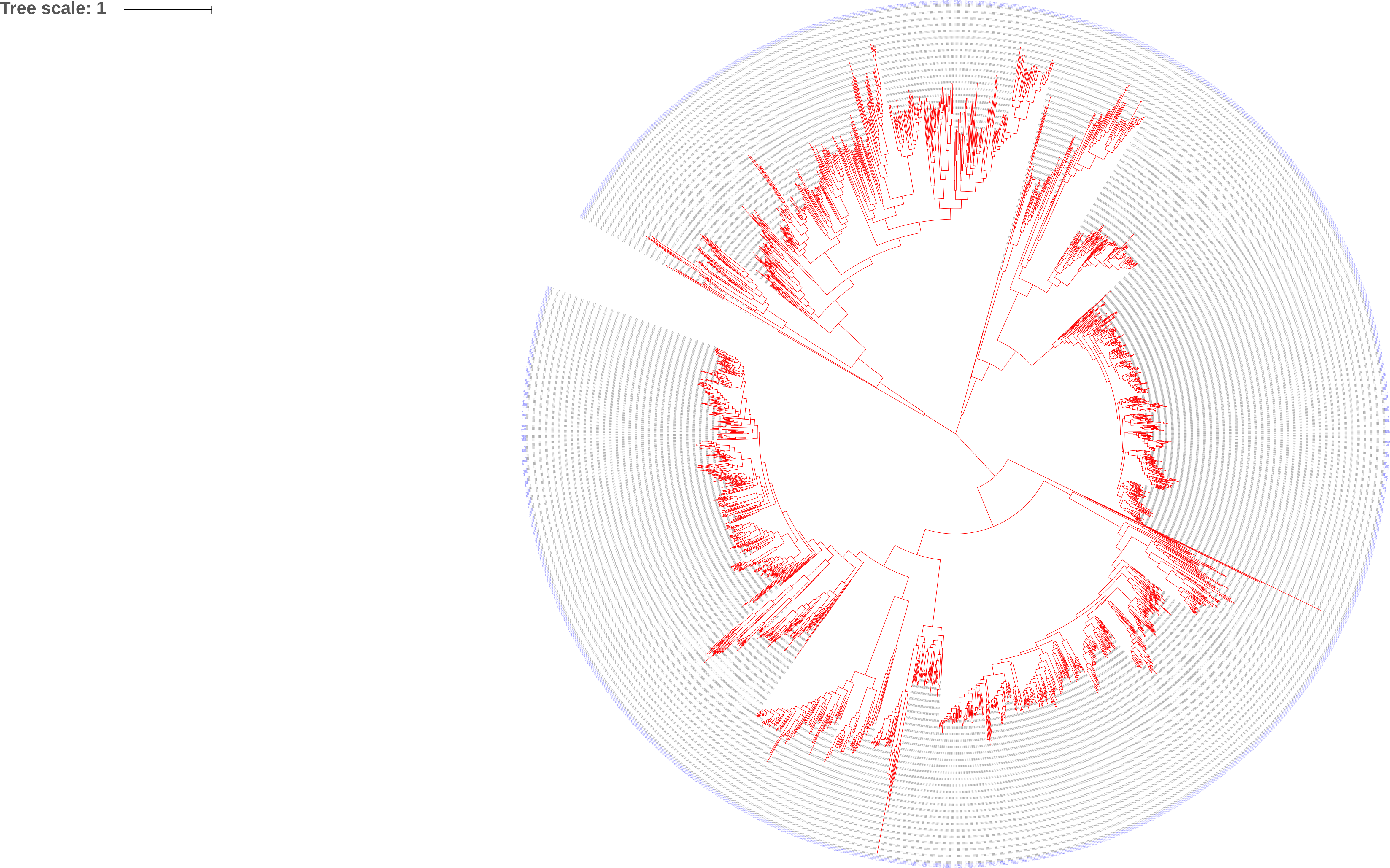

### tRMe.pdf

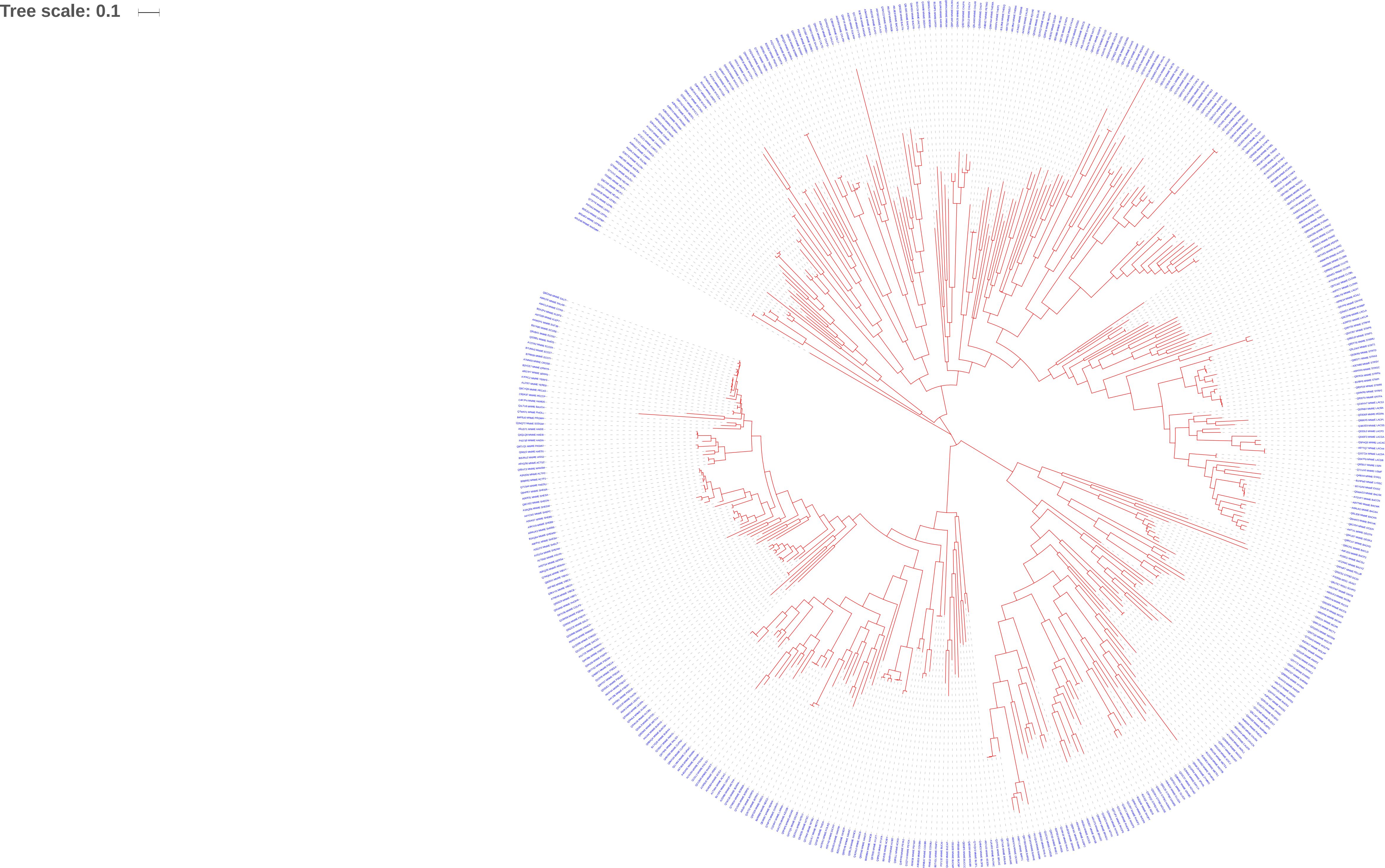
